# Divergence estimation in the presence of incomplete lineage sorting and migration

**DOI:** 10.1101/174342

**Authors:** Graham Jones

**Affiliations:** Department of Biological and Environmental Sciences, University of Gothenburg, Box 461, SE 405 30 Göteborg, Sweden.

## Abstract

This paper focuses on the problem of estimating a species tree from multilocus data in the presence of incomplete lineage sorting and migration. We develop a mathematical model similar to IMa2 (Hey 2010) for the relevant evolutionary processes which allows both the the population size parameters and the migration rates between pairs of species tree branches to be integrated out. We then describe a BEAST2 package DENIM which based on this model, and which uses an approximation to sample from the posterior. The approximation is based on the assumption that migrations are rare, and it only samples from certain regions of the posterior which seem likely given this assumption. The method breaks down if there is a lot of migration. Using simulations, Leaché et al 2014 showed migration causes problems for species tree inference using the multispecies coalescent when migration is present but ignored. We re-analyze this simulated data to explore DENIM’s performance, and demonstrate substantial improvements over *BEAST. We also re-analyze an empirical data set. [isolation-with-migration; incomplete lineage sorting; multispecies coalescent; species tree; phylogenetic analysis; Bayesian; Markov chain Monte Carlo]

---

Speciation is probably gradual in most cases (Martin et al. 2013) and gene flow may continue to occur even between good species. It is difficult to infer these phenomena (Nosil 2008), but some large scale studies have demonstrated them in (for example) butterflies (Martin et al. 2013), birds (Rheindt et al. 2014), grasses (Romaschenko et al. 2014), and big cats (Figueiró et al. 2017). Meanwhile, almost all estimates of species trees to date have been conducted under the assumption that no migration occurs after an instantaneous speciation event. Leaché et al. (2014) simulated a large set of data containing migrations and ILS, and analysed it using *BEAST. They showed that migration causes problems for species tree inference using the multispecies coalescent when migration is present but ignored. In general, migration between sister species rarely caused topological errors, although it resulted in biased estimates of node heights and population sizes. Migration between nonsister species resulted in topological errors, even when the migration rates were small.

This paper describes a a BEAST2 package DENIM for inferring a species tree in the presence of incomplete lineage sorting (ILS) and migration. DENIM allows arbitrary numbers of species, and allows for different (and asymmetric) migration rates between each pair of contemporaneous branches. DENIM is appropriate when the focus is on estimating the species tree when migration rates are small, but it is not suitable when migration rates may be large. As well as inferring the species tree, it is possible to identify which loci are subject to migration, and to infer the time of migration events and the source and destination branches in the species tree.

An alternative approach to gene flow is to model the situation with a species network instead of a tree (for example, Solís-Lemus and Ané (2016); Wen et al. (2016); Zhang et al. (2017)). If hybrid speciation occurs, or populations merge, then a network is essential to model the situation properly. Here we assume that there is a species tree and that any gene flow between pairs of branches will eventually become zero.

‘Migration’ is used to refer to gene flow between species (usually introgression but not restricted to that). We use the term ‘species’ rather than ‘population’ because the method is aimed at situations where gene flow is small. A migration event occurs when an allele comes from a parent from another species. An ‘embedding’ of a gene tree specifies which species tree branch each coalescence belongs to, together with migration events, which specify the times along gene tree branches at which an allele moved between species tree branches, and which species tree branches are involved. We always describe events going back in time from the present, so alleles have parental species to which they ‘go’, and the ‘destination’ branch in the species tree contains part of a gene tree branch at a more ancient time than the ‘source’ branch. This is because coalesences are easier to model this way, and is the same convention as the program IMa2 (Hey and Nielsen 2004, 2007, Hey 2010).

There is no upper limit on the possible number of migration events, and even if this is limited, and we just consider a fixed species tree and a single fixed gene tree, there can still be a huge space of possible embeddings. It is thus difficult to make inferences if the situation modelled in full. In particular it appears very difficult to design and implement MCMC operators capable of sampling efficiently from the full distribution while estimating the species tree. IMa2 requires that the true population phylogeny (equivalent to species tree here) is known. The methods of Tian and Kubatko (2016) and Dalquen et al. (2016) are restricted to 3 species. PHARPL (Jackson et al. 2017) uses a very different type of approximation.

We use a model for migration which is similar to that used by Hey (2010) in IMa2. There are two migration rate parameters for each pair of contemporaneous species tree branches, which means there are 2(*n* − 1)^2^ of them for a species tree with *n* tips (Hey 2010). There are three main differences between DENIM and IMa2. We estimate the species tree instead of assuming it; we integrate out the migration rate parameters; and we use an approximation to simplify sampling from the posterior. This approximation ignores most of the embeddings which seem unlikely given small migration rates, and it restricts the maximum number of migration events per gene tree to be no more than the number of coalescences. If the migration rates is high, some of the ignored embeddings will be quite likely and the approximation will break down. We also integrate out the population size parameters in a similar fashion to Jones (2016).

We first describe the underlying evolutionary model for the coalescent and migration processes, and the particular formulation which allows us to integrate out the population size parameters, and the migration rates. Then we describe the approximation which allows us to sample efficiently from the posterior. The simulated data of Leaché et al. (2014) is re-analysed to evaluate the proposed method, and the pocket gopher data of Belfiore et al. (2008) provides an empirical demonstration of the method.

## The prior density for a gene tree

### Background

Following the introduction of the Kingman coalescent (Kingman 1982), models for coalescence and migration were developed in the 1980s by population geneticists (Hudson et al. 1990). More recent developments include Beerli and Felsenstein (2001), Ewing and Allen (2006), Tian and Kubatko (2016), Dalquen et al. (2016) as well as the work of Hey and Nielsen. The methods of Tian and Kubatko (2016) and Dalquen et al. (2016) can estimate the species tree, but are currently restricted to at most 3 species and 3 sequences per locus.

The underlying evolutionary model we use here is the same as that of Hey (2010), except that the species tree *S* is not assumed known but instead follows a birth-death model. When the species tree *S* is estimated, it is important that ∫ Pr(*G*|*S*)*dG* = 1 for any *S*, where *G* is a gene tree. Here we are assuming that *S* and *G* each have a fixed set of tip labels, and that the tips of *G* are assigned to the tips of *S*. I have not found a clear statement to this effect in the literature, so some explanation seems warranted. Between the node heights of the species tree, we have an *n*-island model for coalescence and migration (Beerli and Felsenstein 2001), where *n* is the current number of species tree branches. This is a continuous time Markov chain. It could be time-inhomogeneous, to allow for population sizes or migration rates to vary continuously with time, although our application here only uses the time-homogeneous case. In order to define the state space of this Markov chain, we need a few preliminaries.

Firstly, each branch in *G* is labeled by the tip labels that descend from the branch. When a coalescence occurs, it should be understood as the merging of two particular labeled gene tree branches. Likewise, when a migration occurs, a particular gene tree branch migrates to a particular species tree branch. Let *L* be the set of tip labels of *G*, and let 𝒫(*L*) be the set of all partitions of *L*. Each partition *P* ∈ 𝒫(*L*) is a set {*L*_1_,…, *L*_*m*_} for some *m* with 1 ≤ *m* ≤ |*L*|, where each *L*_*i*_ is a nonempty subset of *L*, the union of them all is *L*, and they are pairwise disjoint. The subsets *L*_*i*_ are called the ‘blocks’ of the partition. At any time, the set of gene tree branches can be regarded as a member of 𝒫(*L*), and each branch as a block. We will call the periods between node heights of *S*, during which the number of branches is constant, an ‘epoch’. The branches of *S* could be labeled in a similar manner to *G*, but for convenience, we assume they have been labelled with the numbers {1,…, *n*} during the epoch when there are *n* branches, and that branches *n* and *n* − 1 merge to form a branch *n* − 1 in the next epoch.

The state space of the Markov chain during the epoch with *n* branches consists of all possible assignments of all members of 𝒫_*n*_(*L*) to the branches of *S*. Each state is a pair (*P*, *f*) where *P* ∈ 𝒫_*n*_(*L*) and *f* is any map from *P* to {1,…, *n*}, assigning gene tree branches to species tree branches. We use the set theory notation *X*^*Y*^ to denote the set of all maps from set *Y* to set *X*. So we can write the state space 𝒜_*n*_ as

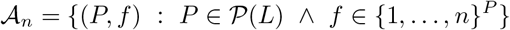

It has size

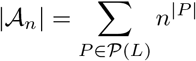

There is an instantaneous rate matrix *Q* of size |𝒜_*n*_| × |𝒜_*n*_|. The off-diagonal rows of *Q* are non-negative, the rows of *Q* sum to zero, and the diagonal entries are less than or equal to zero. In fact all the diagonal entries are strictly negative, except that *Q*_*z*,*z*_ = 0 where *z* is the final state in the root of the species tree, when *n* = 1, and there single gene tree branch. Note that although *Q* is enormous for large |*L*| and *n*, it is sparse, since the number of states which can be reached from a given state by a single migration or coalescence is much smaller than |𝒜_*n*_|. Basic properties of Markov chains (in particular the fact that rows of *Q* sum to zero) ensure that given a starting distribution over states such that

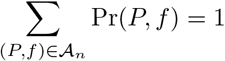
this remains true throughout the process, and in particular just before a merging of species tree branches. At such a merge, the partitions *P* are unchanged, but the state space changes.

Once we are in the root branch of the species tree, the process reduces to the Kingman coalescent, which is a (normalized) density. Consider the case just above the root, where *n* = 2. We have
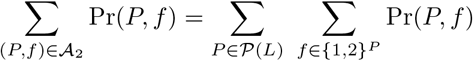

Each *P* consist of blocks *L*_1_,…,*L*_*m*_, and as *f* runs over the maps from *P* to {1, 2}, it runs over exactly those assignments of these blocks to {1, 2} which result in all of them ending up in the root just after the merge. Thus

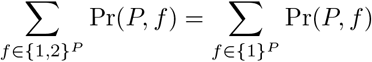

where the left hand side applies just before the merge and the right hand side applies just after the merge. It follows that

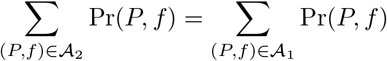

where again the left hand side applies just before and the right hand side applies just after the merge. We can apply a similar argument to merges when *n* > 2 to establish that ∫ Pr(*G*|*S*)*dG* = 1. We will refer to this this evolutionary model as the ‘tree-island model’.

### Integrating out population and migration parameters

Suppose the species tree has s tips. There are 2*s* − 1 species tree branches, including the root branch. Suppose the migration rate from branch *b* to branch *d* is *m*_*bd*_. These migration rates follow the same conventions as Hey and Nielsen (backwards from present, scaled by population size). As in Jones (2016), the population size parameter *θ*_*b*_ for branch *b* is equal to *N*_*b*_*μ*_*b*_, where *N*_*b*_ is the effective population size and *μ*_*b*_ is the mutation rate for the branch. For locus *j*, the effective number of gene copies is obtained from *N*_*b*_ by multiplying by a factor *p*_*j*_ (sometimes called the ‘ploidy’) for gene *j*.

The time (going back from zero at present) is divided into a number of intervals 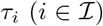 by the times of the events and species tree node heights. The set of species tree branches which exist during the *i*th interval is denoted by 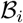, and we set 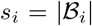. The number of lineages in gene tree *j* which belong to species tree branch *b* during the ith interval is *n*_*jbi*_. The set of intervals which end in a coalesence is 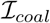, and the set which end in a migration is 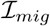. See Figure 1, where 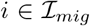, *i* + 1 is in neither, and 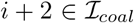. The rate at which the next event occurs is (*κ*_*i*_ + *μ*_*i*_) where

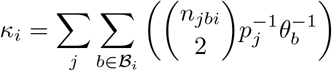

is the total rate for coalescent events and

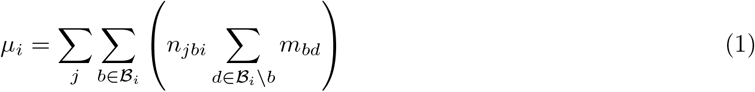
is the total rate for migration events. Here we are summing the nonzero off-diagonal elements of a row of *Q* in order to find *Q*_*x*,*x*_ = − (*κ*_*i*_ + *μ*_*i*_) for the current state *x*. We then need *Q*_*x*,*y*_, where *y* is the next state. If it is a coalescence, 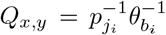 where *j*_*i*_ is the gene tree containing the coalescence, and *b*_*i*_ is the species tree branch in which it occurs. If it is a migration, 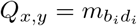, where *b*_*i*_ and *d*_*i*_ are the source and destination branches.

**Figure 1:**
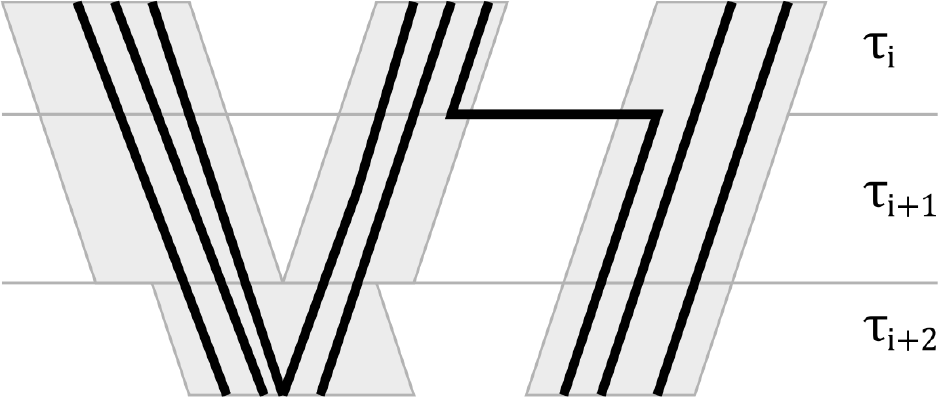
Three time steps. The first ends in a migration, the second with a species tree node, and the third with a coalescence.

Denoting the set of migration rates by *M* and the set of population size parameters by Θ, we have the probability density for a gene tree:

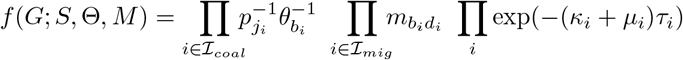

This can be factored into a coalescence part and a migration part. Then, our aim is to rearrange the terms in the coalescence part so that it is a product over species tree branches, and the rearrange the terms in the migration part so that it is a product over ordered pairs of species tree branches. The result will be a product of terms, in which each term contains one population size parameter *θ*_*b*_ or one migration parameter *m*_*bd*_. This enables us to integrate out these parameters if suitable priors are assumed. We put

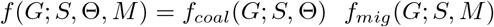

where

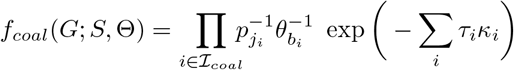

and

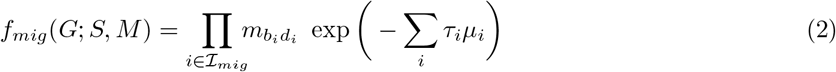

First we deal with *f*_*coal*_. We have

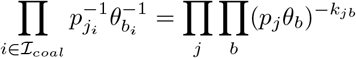

where *k*_*jb*_ is the number of coalescences in gene tree *j* in branch *b*. Next

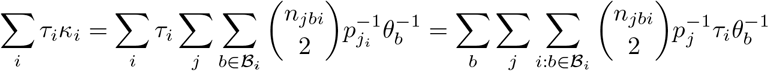

so

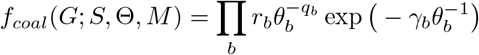
where

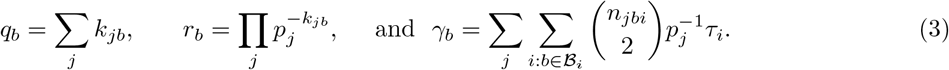

As written, there are time intervals in branch *b* events during which no change occurs in branch *b*. For the computation of *γ*_*b*_, we only need to take into account coalescences within branch *b* and migratitotal rate for migration events. Hereons in and out of branch *b*, since between these events, *n*_*jbi*_ is constant. Equation (3) is now of the same form as equation (2) of Jones (2016). The only difference is that *γ*_*b*_ accounts for migrations in and out of branch *b*. This means the population size parameters can be integrated out as in Jones (2016).

Now we turn to the migration part *f*_*mig*_. Let 𝒪 be the set of contemporaneous pairs of branches in *S*. We have

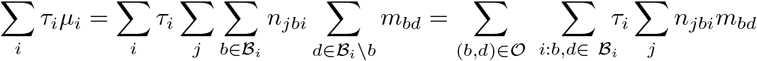

Thus

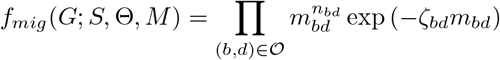

where *n*_*bd*_ is the total number of migrations from *b* to *d* and

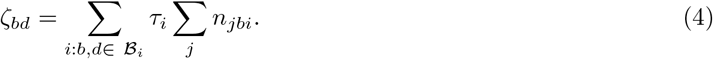

The term ζ_*bd*_ can be interpreted as the total intensity of migrations from *b* to *d* during the time in which both branches *b* and *d* exist. If we assume that *m*_*bd*_ ~𝒢(*α*_*bd*_, *β*_*bd*_) for all *b*, *d* where 𝒢 is the gamma distribution, then we get a contribution to the posterior which is

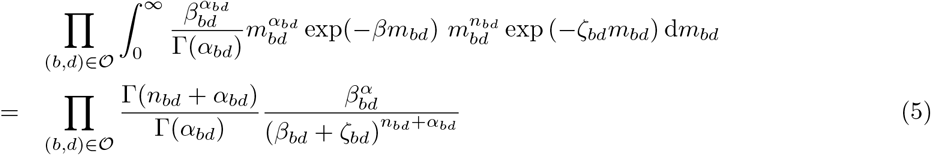

Equations (4) and (5) provide the information needed to implement the calculation for the migration part of the posterior. We have allowed each ordered pair of contemporaneous branches (*b*, *d*) to have a different prior. For example, we can represent the prior expectation that migration rates are lower between more distantly related branches. We will call this model, where migration rates are independent, the ‘flexible’ model.

The calculation in equation (4) is slow when the number of tips in the species tree is large. A much simpler model is to assume that *m*_*bd*_ is the same value *m* for all *b*, *d*. In this case, equation (1) reduces to 
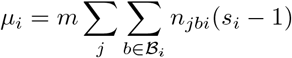

The double sum is equal to the total number of gene tree lineages *N*_*i*_ during time interval *i*. Then we have

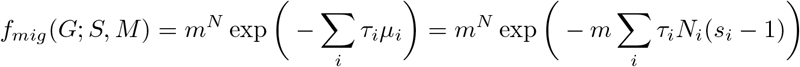
where *N* is the total number of migrations. The parameter *m* can be integrated out. We will call this model the ‘simple’ model.

## How the gene tree is embedded

This section describes the embedding parameters, and how they are used to embed the gene trees. We restrict the embeddings by ignoring ones which are unlikely when the migration rates are small enough. The hope is that we will still explore a region of parameter space which includes most of the probability content.

Embeddings are restricted by applying the following rules:

1. there is at most one migration in a single gene tree branch
2. at most one of the child branches of a gene tree node contains a migration
3. migrations are only used when choosing the embedding if needed (in a sense described below)

We call a pair of child branches of a gene tree node a **sister-pair**. The embedding parameters *E*_*j*_ consist of two values *ξ*_*ji*_, *η*_*ji*_ ∈ [0,1] for each internal node *i* of the *j*th gene tree. See Figure 2. The parameter *ξ*_*ji*_ determines where along a sister-pair a migration may occur, if a migration is needed in the embedding. Thus it determines which child branch of node *i* is capable of migrating, as well as the time of the migration if there is one. All the nodes in the species tree have their children labeled as ‘left’ and ‘right’, so that [0,1] can be mapped unambiguously onto the sister-pair. The other parameter *η*_*ji*_ specifies which of the node’s child branches to use when choosing a destination species branch for an migration. If the migration is between sister branches of the species tree, there is only one choice for the destination. It may happen that the sister branch is too ancient, in which case several destination species branches are possible. The possible destination branches are found, and *η*_*ji*_ is used to choose between them by dividing the interval [0,1] equally into the appropriate number of parts.

**Figure 2:**
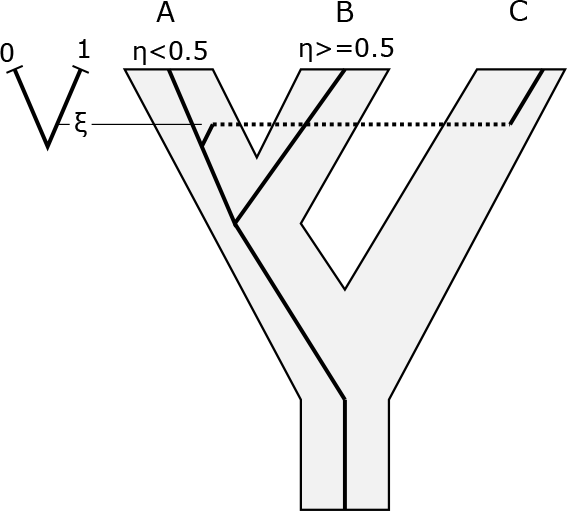
This shows *ξ*_*ji*_ and *η*_*ji*_ for a one gene tree and one node (with subscripts dropped). There is a migration from C into A. The parameter *ξ* determines how far along the sister-pair the migration occurs, and *η* determines whether the destination branch is A or B.

The parameters *ξ*_*ji*_ and *η*_*ji*_ are changed by operators during the MCMC algorithm, regardless of whether or not they are being used to embed a gene tree. This is a simpler alternative to implementing rjMCMC operators which account for changes in dimension. The prior Pr(*E*_*j*_) for *E*_*j*_ is are independent uniform distributions on [0,1] for each *ξ*_*ji*_ and *η*_*ji*_.

The first rule above is straightforward. The definition of *ξ*_*ji*_ enforces the second rule. The third rule is applied recursively from the tips. Suppose *x* is the *i*th node of the *j*th gene tree, and suppose both child nodes of *x* have been assigned to branches in the species tree. If it is possible to assign *x* to a species tree branch without an migration in either child branch of *x*, then this is done. Otherwise *x* is assigned using the species tree branch to which its non-migrating child has been assigned: it will be the same branch, or an ancestor of that branch, depending on the height of *x*. The height of the migration is fixed by *ξ*_*ji*_. The migrating child branch of *x* starts in the species tree branch that the migrating child has been assigned. It stays in this branch, or an ancestor of it, until the migration height. It will then migrate to the same species tree branch as *x*, or a descendant of it. If there is more than one descendant of the species tree branch of *x* at this height, values from *η*_*j*_ are used to choose one.

### Properties of the embedding scheme

Different embeddings of the same gene tree in the same species tree are obtained by changing *ξ*_*ji*_ and *η*_*ji*_ during the MCMC sampling. Figure 3 shows some examples. Case (a) is simple. No migrations are needed to embed the gene tree, so embeddings with one or more migrations are ignored. Case (b) requires one migration, and an embedding with two migrations in the same branch is ignored. Case (c) requires two migrations. The embedding on the left is ignored since it has two sister branches with migrations. The embedding on the right is one of four embeddings that is considered.

Proposition. *Given any set of particular values for *ξ* and *η*, and the rules above, any gene tree can be embedded in any species tree. For any *G*_*j*_ and *S*, the set of embeddings as *ξ* and *η* vary include at least one with a minimal number of migrations.*

Proof: The first claim is straightforward, using recursion starting at the tips, and following the description above (for applying the third rule).

For second claim, suppose it is false and consider the set *M* of minimal embeddings (those with a minimal number of migrating branches). Call a node both of whose child branches migrate a ‘double node’. Thus every member of *M* has at least one double node. Now restrict attention to the subset *M*\* of *M* of embeddings which have as few as possible double nodes. Finally, choose an embedding *B* from *M*\* so that a double node *x* is as near to the root as possible.

If *x* is the root, it can be moved into the same branch as one of its children, or an ancestor of that branch, and one migration can be removed, contradicting the definition of *M*. If *x* is not the root, it can be again moved into the same branch as one of its children, but now the branch between *x* and its parent may need to become migrating. If the sister branch to *x* is already migrating, we have an embedding with the same number of migrations, but a double node closer to the root than *x*, contradicting the definition of *B*. If the sister branch to *x* is not migrating, we have an embedding with fewer double nodes than *B*, contradicting the definition of *M*\*. End of proof.

Thus the scheme ensures that at least one minimal embedding is considered. However, the method does not consider every embedding which has a minimal number of migrations (e.g., Figure 3c). Also note that some embeddings which are considered are not minimal. For example, consider a species tree (A,B) and a gene tree ((a1,b1),b2) with three tips a1, b1, and b2, where a1 belongs to species A and the others to B. Suppose the species tree has greater height than the gene tree. For some values of the continuous parameters *η* and *ξ*, DENIM will assign the coalescence (a1,b1) to A, which means two migrations will be used to embed the gene tree. With other values of *η* and *ξ* the gene tree will be embedded with only one migration.

**Figure 3:**
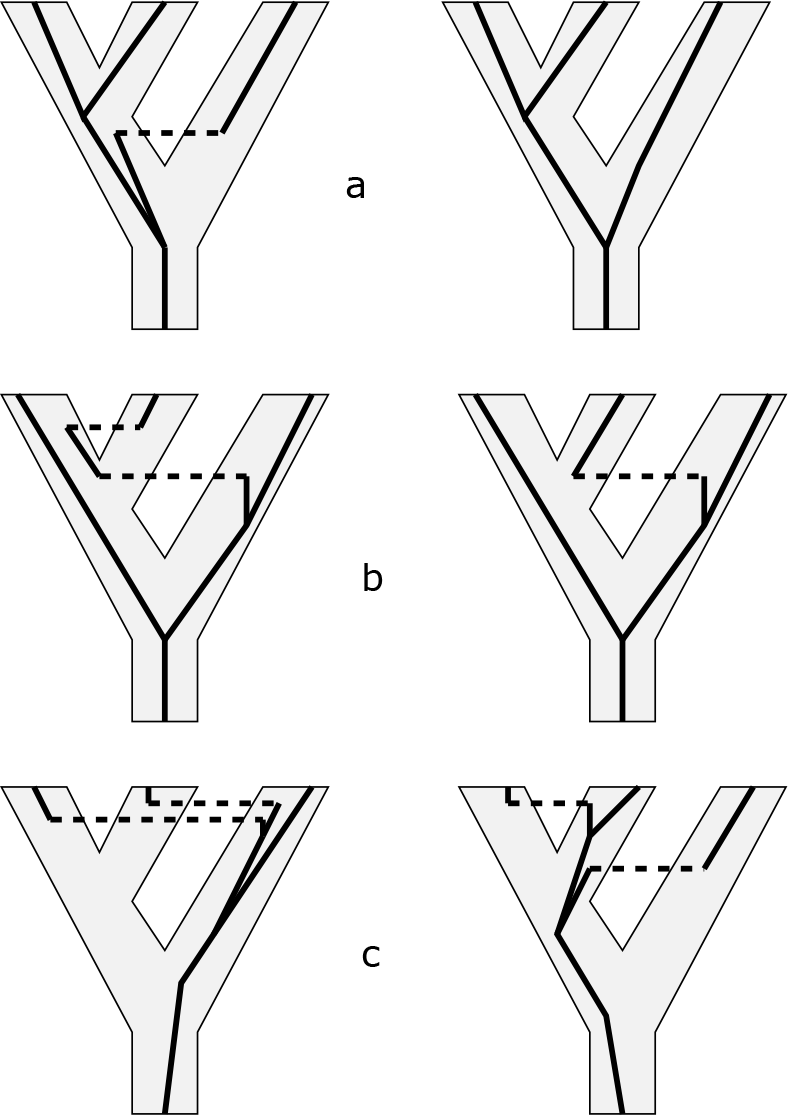
Some examples of embeddings for three gene trees in a, b, c. On the left are embeddings that are ignored. On the right are embeddings which are considered.

## Implementation notes

For a standard multispecies coalescent analysis, operators which change the species tree and the gene trees in a coordinated way are beneficial (Jones 2016; Ogilvie et al. 2017). These operators rely on, and preserve, compatibility between the species tree and gene trees under the multispecies coalescent. In the presence of migration, any gene tree is compatible with any species tree, and unfortunately these coordinated operators cannot be used as they are. In the current implementation, DENIM uses the standard tree operators implemented in BEAST2 for the species tree and the gene trees. A couple of simple MCMC operators were implemented for the embedding parameters. As noted above, they are changed by operators regardless of whether or not they are being used to embed a gene tree. In general, applying MCMC operators to unused parameters could be very inefficient, and rjMCMC would be preferable, but here the operators for *ξ* and *η* are very fast.

DENIM is implemented in the BEAST2 framework, and so benefits from the flexible site models, substitution models, and others available in BEAST2. An analysis can be set up using the graphical interface Beauti. When using the flexible model, different priors can be used for different pairs of species tree branches. DENIM provides two schemes for specifying these priors. In the first scheme, the prior mean for the migration rate between branches depends on how closely related the species are, with the prior mean decreasing as the number of branches between them increases. The rate at which the prior mean decreases is controlled by a user-defined parameter called RelatednessFactor. In the second, the migration rate decays with the time between the most recent common ancestor of the two branches and the midpoint of the interval during which both branches exist. The decay is controlled by a user-defined parameter called MigrationDecayScale. The details are described in the manual supplied with DENIM.

DENIM puts some annotations into the species trees sampled during the MCMC process, which give more detail about the migrations. These can be analyzed using the command line program `MigrationAnalyser.jar` which can be downloaded from http://indriid.com/software.html.

## Tests using simulated data

The method was tested using the simulated data from Leaché et al. (2014), and with no data. Version 0.3.0 of DENIM was used for these tests. The simulated sequences and some other files are available from http://indriid.com/workingnotes2017.html. For all these tests, an exponential prior for the migration rate was used; the mean of this prior was varied in some tests. No use was made of the options to use different prior means for different pairs of species tree branches, so within each analysis, the same prior is used for all pairs. This choice of prior for the migration rates can be made in Beauti by setting MigrationDecayScale to a negative value, and RelatednessFactor to 1.0.

The MCcoal program (Yang 2015) was used to generate the sequence files. The MCcoal program was extended slightly to make it produce information about each individual migration in the simulations, instead of the normal summary information. This enables tests of how well DENIM can infer the existence of migration events per locus.

The MCcoal control files were the same as those of Leaché et al. (2014), except that in the 10-species scenarios, they were augmented by adding scenarios for a migration rate of 0.01, making 25 migration patterns in total. In Leaché et al. (2014), only the two highest rates 0.1 and 1.0 were used in the 10-species scenarios, but preliminary results showed that DENIM generally breaks down between 0.01 and 0.1, so the 0.01 rate is an interesting one for DENIM.

### Prior only

We tested DENIM by running it with no data. The scenarios used all had 6 species. The prior on the species tree was a pure birth (Yule) model with growth rate 100. The expected height of the species tree can be calculated as 0.0145 (?). The number of individuals *i* was 3 or 9 per species, the number *g* of loci was 3 or 9. The population scaling parameter in DENIM was set 0.0005, 0.005, 0.05, producing small, medium, and large amounts of ILS. Finally we tried both the flexible and simple models, resulting in a total of 2 × 2 × 2 × 3 × 2 = 48 scenarios. The analyses were run for 30M (*i* = 3, *g* = 3), 90M (*i* = 3, *g* = 9 and *i* = 9, *g* =3), 300M (*i* = 9, *g* = 9) generations. These long runs proved necessary to obtain convergence.

### 4-species scenarios

The simulated data of Leaché et al. (2014) was used with the simple model. An prior mean of 0.005 was used for the migration rate. All 100 replicates were used. The chain length was 10M, states were logged every 5000 generations, and burnin was set to 20%, or 400 out of 2000 states.

The settings for site and clock models were similar to those used by Leaché et al. (2014). DENIM uses a different population model to *BEAST, so this is somewhat different. Site models were linked. A GTR model of substitution was used, with base frequencies equal. The clock models were strict but unlinked. The first locus had clock rate 1, and the others were estimated. The Yule (pure birth) model was used for the species tree. The priors were set as follows. Substitution rates relative to rateCT: Gamma(0.05,20). Relative clock rates: lognormal(0,1). Growth rate for the species tree: lognormal(5,2). PopPriorScale: lognormal(-5,2).

Further experiments were conducted using the first 25 replicates. The settings were the same as above, except that the flexible model was used, and the prior means for the migration rate were varied: the set of values (0.00125, 0.005, 0.02, 0.08) were used. The flexible model was used since the experiments with priors suggest that this model is likely to be better with high migration rates in the prior.

### 10-species scenarios

There were some convergence problems, so the chain length was increased to 20M, with burnin at 20%, or 800 out of 4000 states. Only the first 50 replicates were analyzed. Other settings were as for the 4-species case, using the simple model and an prior mean of 0.005 for the migration rate.

## Results on simulated data

### Evaluation measures

We use three measures for assessing accuracy. One measure is the probability coverage. This is the proportion of replicates where the true species tree topology is in the 95% credible set. This measure is used by Leaché et al. (2014) so a direct comparison can be made between *BEAST and DENIM. However, this measure does not take into account errors in estimated node heights.

The second measure is based on the branch score of Kuhner and Felsenstein (1994), adapted for rooted trees. It accounts for differences in topology and branch lengths. The entire posterior is evaluated by finding the mean distance between the MCMC samples of the species tree and the true tree. We use this as our main overall measure of accuracy for comparing different settings within DENIM.

The third measure is aimed at evaluating how well DENIM can identify which loci are migrating. For each locus, DENIM outputs a statistic which counts the number of migrations in the MCMC sample. The posterior mean of this count can be compared to the true number. Here we only report the posterior means for these counts in the two cases in which the true number is either zero or greater than zero, that is, that migration is either absent or present in the simulated data for a locus. DENIM produces a lot more detail about migrations but it is difficult to summarize for a large number of replicates and scenarios.

### Prior only

Figure 4 shows some results from using DENIM with no data. In the upper graph, the values should all be close to zero, corresponding to a ratio (estimated height)/(true height) near 1. Instead, some values are much larger, corresponding to a ratio of 10 or more. The large estimates of the species tree height are accompanied by large migration counts. Thus, with no data and a large number of loci and individuals, the method breaks down even with small prior mean for the migration rate. The flexible model behaves better than the simple model.

**Figure 4:**
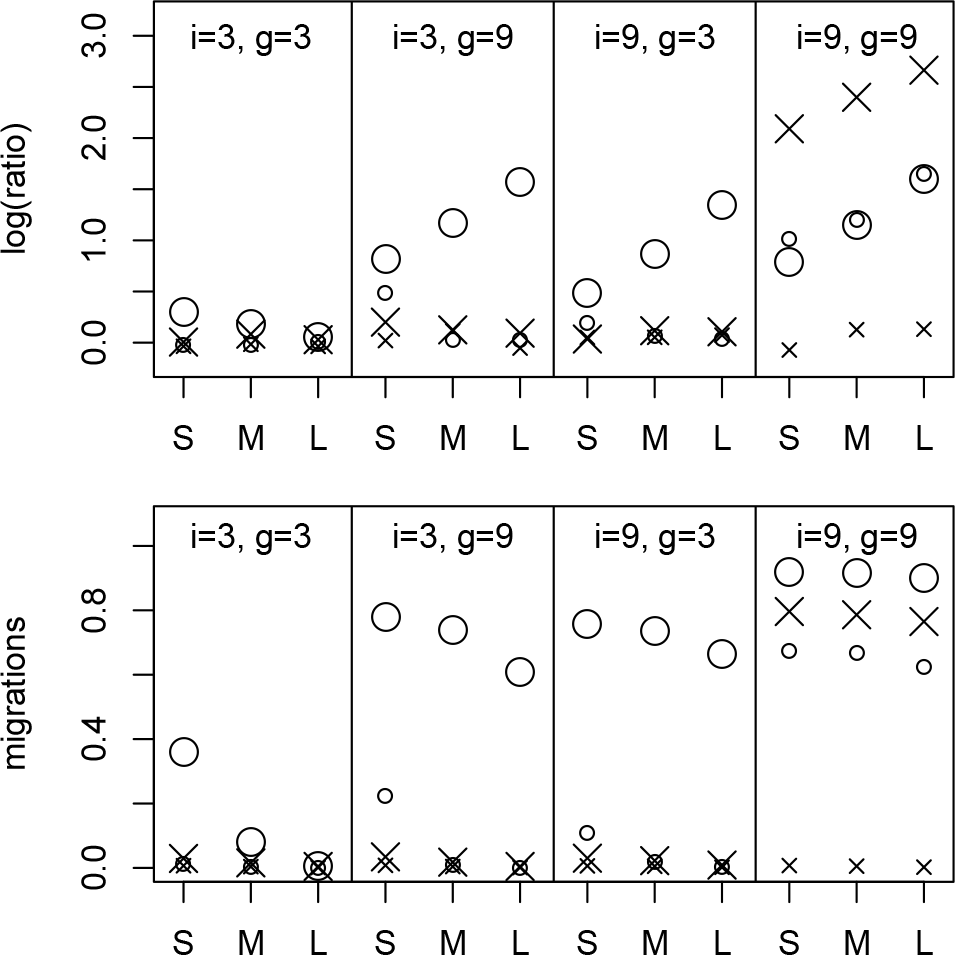
Results with no data. The *y*-axis in the top graph is the log-ratio of the estimated species tree height to the true species tree height. i is the number of individuals per species, and g is the number of loci. S, M, and L represent small, medium, and large amounts of ILS. The *y*-axis in the bottom graph is the proportion of migrations per coalescence (or per sister-pair). Circles represent the simple model, crosses represent the flexible model. Large symbols correspond to a prior mean for the migration rate of 0.005, small ones to a prior mean of 0.001.

It is not understood why the approximation used by DENIM leads to this behavior, but the following may offer some insight. Consider the case of a species tree *S* and a gene tree *G* with just two tips each. We can divide the parameter space into regions corresponding to 0,1,2,… migrations. If there are none, DENIM will sample correctly from the prior. Given the small prior mean for the migration rate, the regions corresponding to 2 or more migrations are not significant. The main bias arises where there is one migration. For height(*S*) > height(*G*), a migration is needed, and DENIM considers this case. If height(*S*) < height(*G*), DENIM only considers the possibility of no migration. The region of parameter space with height(*S*) < height(*G*) is most affected by the approximation used by DENIM, so that small values of height(*S*) are discriminated against. The general effect of this sort of bias apparently gets worse as the number of loci and individuals increases. It is also worth noting that as the species tree becomes taller, the expected number of migrations increases, so there some ‘feedback’ once things have started to go wrong.

### Simulated data: Effective sample sizes

Effective sample sizes (ESSs), as estimated by CODA (Plummer et al. 2006), for the posterior were generally above 200. In the 4-species scenarios, of a total of 1700 replicates, 89 were in the range 100-200, 14 were in the range 50-100, and one (`B-C_0p1` replicate 37) was only 15. In this case, the posterior and the migration counts jumped upwards at around 4M generations. In the 4-species scenarios, of a total of 1250 replicates, 213 were in the range 100-200, 9 were in the range 50-100, and 3 were in the range 25-50. Clearly longer runs would be better, but these were already time-consuming (about 5 weeks on a desktop computer with 4 cores). There was no obvious correlation between low ESS and low accuracy.

The species tree root height often had the worst ESS among all parameters. It is not clear why this should be. The root height was generally estimated fairly accurately; and the operators affecting it appeared to be working satisfactorily. Overall, DENIM is slower than *BEAST (v1), but not hugely so.

### Simulated data: Coverage

Tables 1 and 2 show the coverage probability for all scenarios. These are included for direct comparison with Leaché et al. (2014). In general, DENIM performs better than *BEAST in the cases of paraphyly and the migrations at time zero. In two scenarios among the 10-species set, where there is considerable migration between sister species, DENIM is substantially worse. Otherwise, the performance is similar.

**Table 1:**
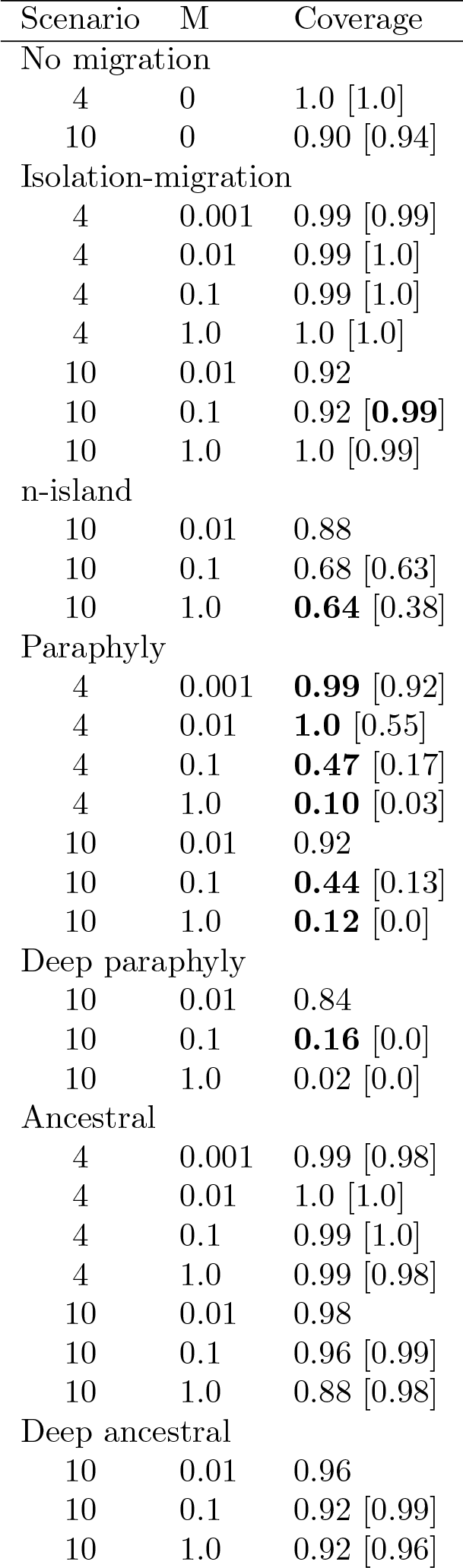
Coverage for scenarios with continuous migration. Values for *BEAST from Leaché et al. (2014) are shown in square brackets. Where the programs produce results which are substantially different, the better result is in bold.

**Table 2:** Coverage for scenarios with continuous migration. Values for *BEAST from Leaché et al. (2014) are shown in square brackets. Where the programs produce results which are substantially different, the better result is in bold.

| Scenario | M | Coverage |
| --- | --- | --- |
| No migration |  |  |
| 4 | 0 | 1.0 [1.0] |
| 10 | 0 | 0.90 [0.94] |
| Single migrant |  |  |
| 4 | Sister species | 1.0 [1.0] |
| 4 | Non-sister species | **0.98** [0.09] |
| 10 | Sister species | 0.86 [**0.98**] |
| 10 | Non-sister species | **0.88** [0.07] |
| Deep single migrant |  |  |
| 10 | Non-sister species | **0.88** [0.0] |
| Single locus introgression |  |  |
| 4 | Sister species | 1.0 [0.99] |
| 4 | Non-sister species | **0.98** [0.37] |
| 10 | Sister species | 0.96 [0.99] |
| 10 | Non-sister species | **0.88** [0.28] |
| Deep single locus introgression |  |  |
| 10 | Non-sister species | **0.88** [0.0] |

Table 3 shows the coverage probability for the 4-species scenarios as the settings in DENIM are varied. With this data, there is little difference between the simple and flexible models. As the prior mean is increased, the results for paraphyletic migration with rates 0.1 and 1.0 improve, while other scenarios become slightly worse. With the smallest prior mean of 0.00125, the result for the scenario with a single migrant between non-sisters becomes worse.

**Table 3:**
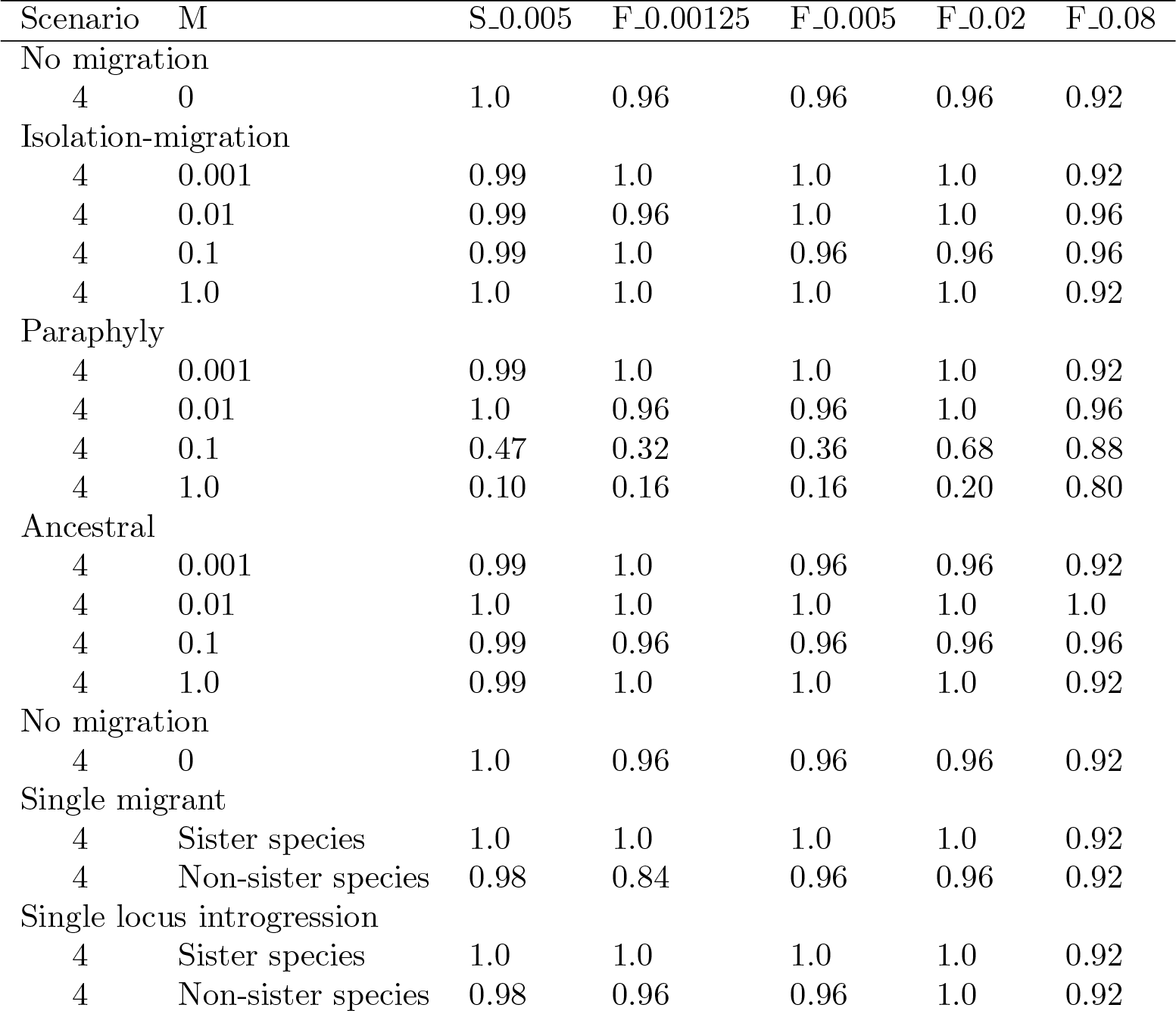
Coverage for 4-species scenarios, and different settings in DENIM. S_0.005 stands for the simple model with prior mean 0.005 on the migration rate, F_0.00125 for the flexible one with prior mean 0.00125, and so on.

| Scenario | M | S_0.005 | F_0.00125 | F_0.005 | F_0.02 | F_0.08 |
| --- | --- | --- | --- | --- | --- | --- |
| No migration |  |  |  |  |  |  |
| 4 | 0 | 1.0 | 0.96 | 0.96 | 0.96 | 0.92 |
| Isolation-migration |  |  |  |  |  |  |
| 4 | 0.001 | 0.99 | 1.0 | 1.0 | 1.0 | 0.92 |
| 4 | 0.01 | 0.99 | 0.96 | 1.0 | 1.0 | 0.96 |
| 4 | 0.1 | 0.99 | 1.0 | 0.96 | 0.96 | 0.96 |
| 4 | 1.0 | 1.0 | 1.0 | 1.0 | 1.0 | 0.92 |
| Paraphyly |  |  |  |  |  |  |
| 4 | 0.001 | 0.99 | 1.0 | 1.0 | 1.0 | 0.92 |
| 4 | 0.01 | 1.0 | 0.96 | 0.96 | 1.0 | 0.96 |
| 4 | 0.1 | 0.47 | 0.32 | 0.36 | 0.68 | 0.88 |
| 4 | 1.0 | 0.10 | 0.16 | 0.16 | 0.20 | 0.80 |
| Ancestral |  |  |  |  |  |  |
| 4 | 0.001 | 0.99 | 1.0 | 0.96 | 0.96 | 0.92 |
| 4 | 0.01 | 1.0 | 1.0 | 1.0 | 1.0 | 1.0 |
| 4 | 0.1 | 0.99 | 0.96 | 0.96 | 0.96 | 0.96 |
| 4 | 1.0 | 0.99 | 1.0 | 1.0 | 1.0 | 0.92 |
| No migration |  |  |  |  |  |  |
| 4 | 0 | 1.0 | 0.96 | 0.96 | 0.96 | 0.92 |
| Single migrant |  |  |  |  |  |  |
| 4 | Sister species | 1.0 | 1.0 | 1.0 | 1.0 | 0.92 |
| 4 | Non-sister species | 0.98 | 0.84 | 0.96 | 0.96 | 0.92 |
| Single locus introgression |  |  |  |  |  |  |
| 4 | Sister species | 1.0 | 1.0 | 1.0 | 1.0 | 0.92 |
| 4 | Non-sister species | 0.98 | 0.96 | 0.96 | 1.0 | 0.92 |

### Simulated data: Branch scores

Figures 5 and 6 show the branch scores for the 4 and 10 species scenarios. The general picture is that DENIM produces good results for the two smallest migration rates 0.001 and 0.01, and for the migrations at time zero, but breaks down at higher rates.

**Figure 5:**
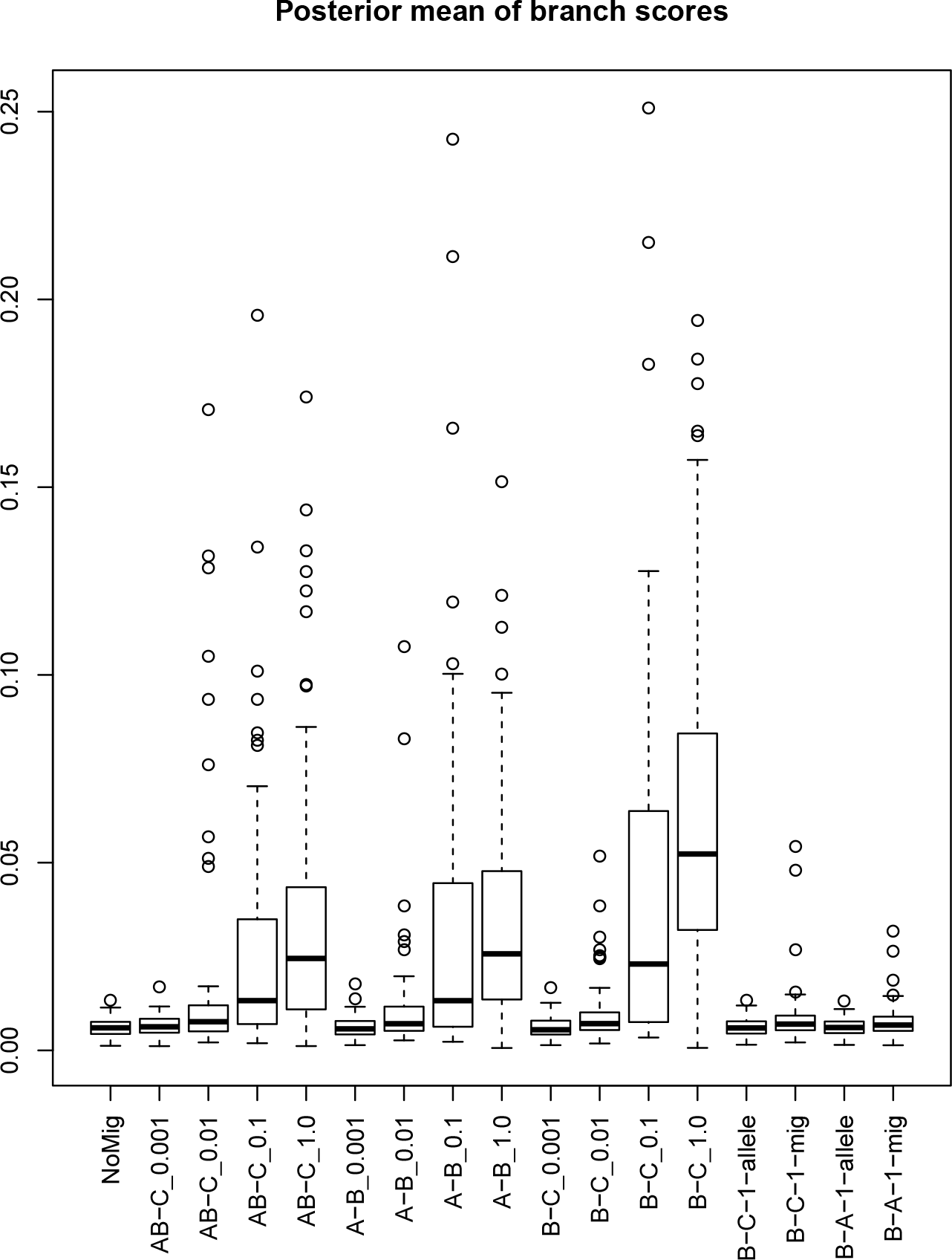
Branch scores for the 4-species scenarios, based on 100 replicates. “NoMig” is the scenario with no migration. The other names describe the pairs of species tree branches which have migration, followed by the migration rate, or “allele” meaning a single locus introgression, or “mig”, meaning a single migrant. The simple model was used with prior mean of 0.005.

**Figure 6:**
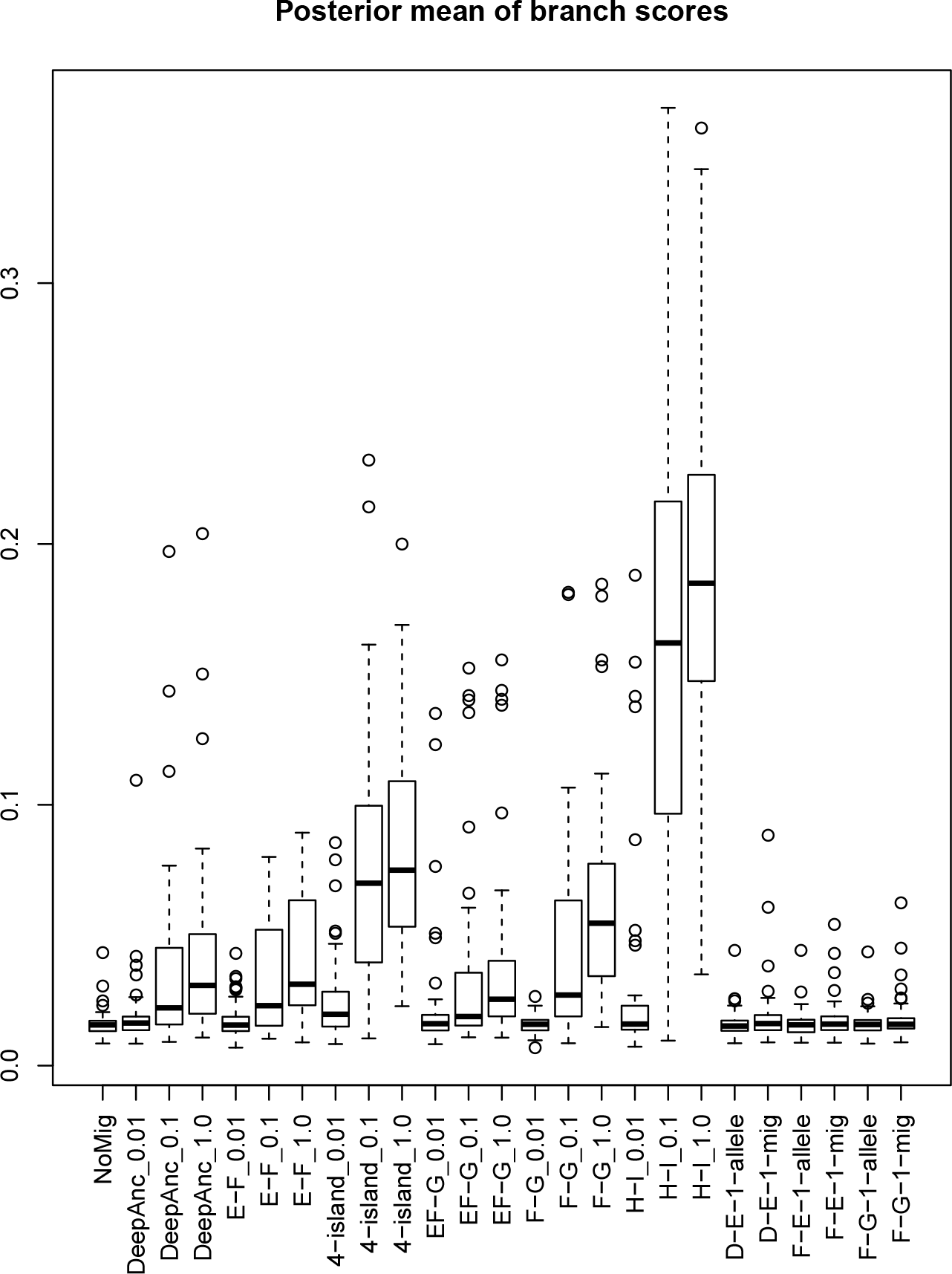
Branch scores for the 10-species scenarios, based on 50 replicates. “NoMig” is the scenario with no migration. “4-island” means there is migration between 6 pairs of branches (E and F, E and G, E and H, F and G, F and H, G and H). “DeepAnc” is the deep ancestral scenario, with migration between ABCD and EFGH. The other names describe the pairs of species tree branches which have migration, followed by the migration rate, or “allele”, meaning a single locus introgression, or “mig”, meaning a single migrant. The simple model was used with prior mean of 0.005.

Supplementary figures 10, 11, 12, 13 show results for the 4-species scenarios with the flexible model, and different prior means 0.00125, 0.005, 0.02, 0.08. The results do not vary greatly with the choice of prior, and follow a similar pattern to the coverage results. All but the high-rate paraphyletic scenarios become worse with a prior mean of 0.08.

### Simulated data: Migration detection

Figures 5 and 6 show the ability of DENIM to infer the existence of migration. The general picture is that DENIM only does this well when there migration rate is low and between non-sister species. DENIM usually ‘explains’ migration between sister species by squashing the species tree (like *BEAST). An exception to the general picture is where there is a single migrant at time zero between sister species (B-A-1-mig in Figure 5, F-E-1-mig in Figure 6) where the migration is usually detected. Supplementary figures 14, 15, 16, 17 show results for the 4-species scenarios with the flexible model, and different prior means 0.00125, 0.005, 0.02, 0.08.

Generally speaking, DENIM identifies loci with migrations which result in an incompatibility with the species tree. Some migrations do not cause incompatibility, because (going back in time) they do not coalesce with another lineage until the species tree branches have merged; or a lineage may migrate, then migrate back again before coalescing, and so on. These are generally missed by DENIM.

## Results on empirical data

We re-analyzed the pocket gopher data of Belfiore et al. (2008). We used the HKY substitution model, linked site models, estimated relative clock rate for all loci except the first, and a strict clock. The results here use the simple model for migration, with an exponential prior with mean 0.001.

This data was also analyzed in Heled and Drummond (2010), the paper which introduced *BEAST. In the *BEAST analysis, the outgroup species *Orthogeomys heterodus* was misplaced (their Figure 8a), and the authors comment that “The tendency to place the outgroup incorrectly appears to be caused by just one gene” namely TBO29. The tree from the DENIM analysis is shown in Figure 9. The outgroup is correctly placed, and it is very similar to the *BEAST result with ingroup monophyly enforced (their Figure 8b). The DENIM tree is somewhat shorter, perhaps due to a different site model or population model. A migration was inferred between *Orthogeomys heterodus* and the (*T. bottae, Thomomys townsendi, Thomomys umbrinus*) clade, the same clades that *BEAST grouped together. This migration was present in about 95% of the MCMC samples. The other migrations that appear in the posterior samples have much lower posterior probabilities. The next migrations that DENIM analysis suggests (at about 24%) are very recent ones, both ways, of TBO47 between *T. bottae* and *T. umbrinus*. This pair is followed (at about 18%) by a very recent one of TBO64 *T. talpoides* to *T. idahoensis* (going back in time).

**Figure 7:**
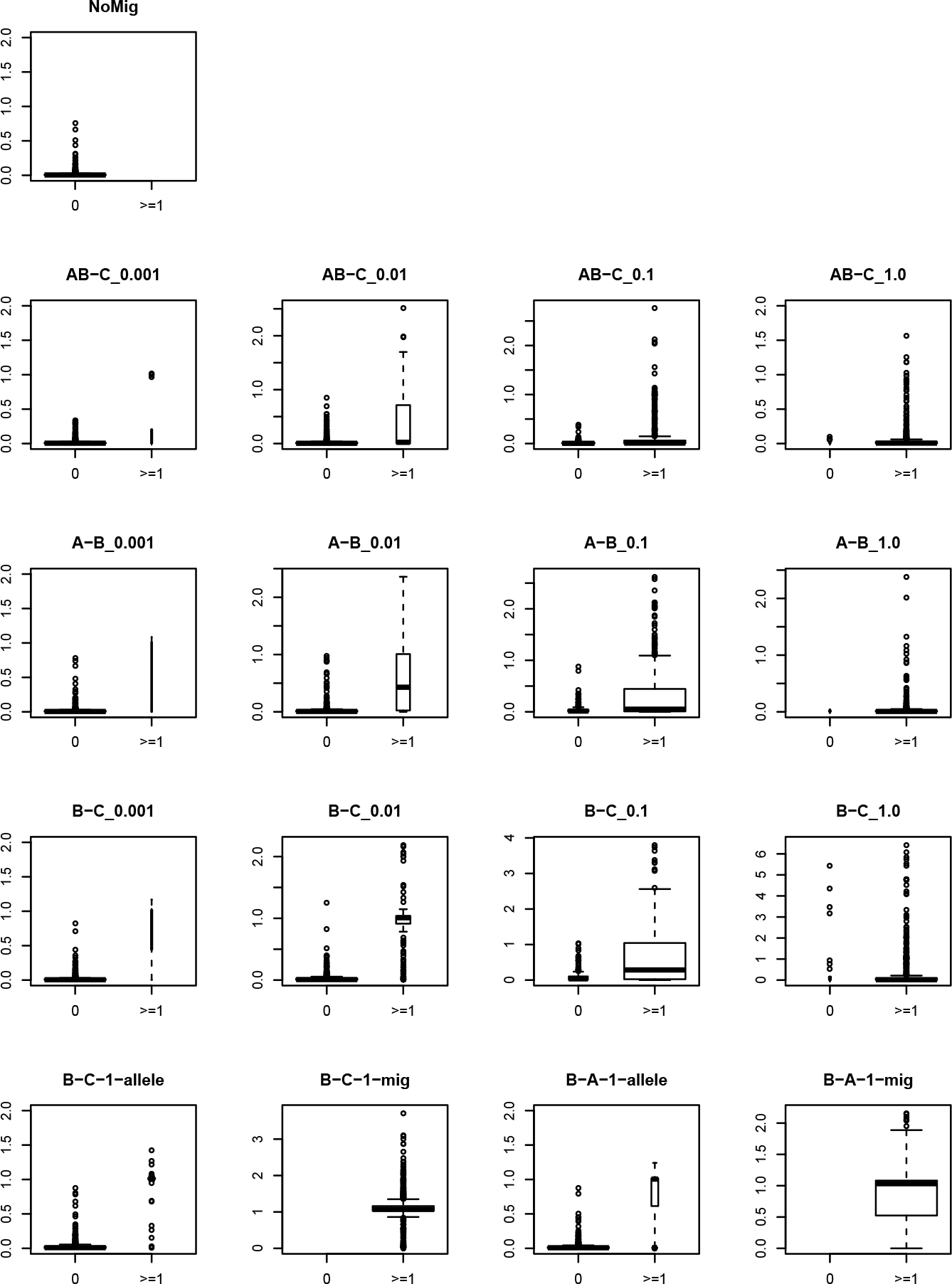
Migration detection for the 4-species scenarios. Each boxplot show the posterior mean count of migrations for the two cases that migration is present or absent in a locus in the simulated data. The width of the boxes is proportional to the number of cases.

**Figure 8:**
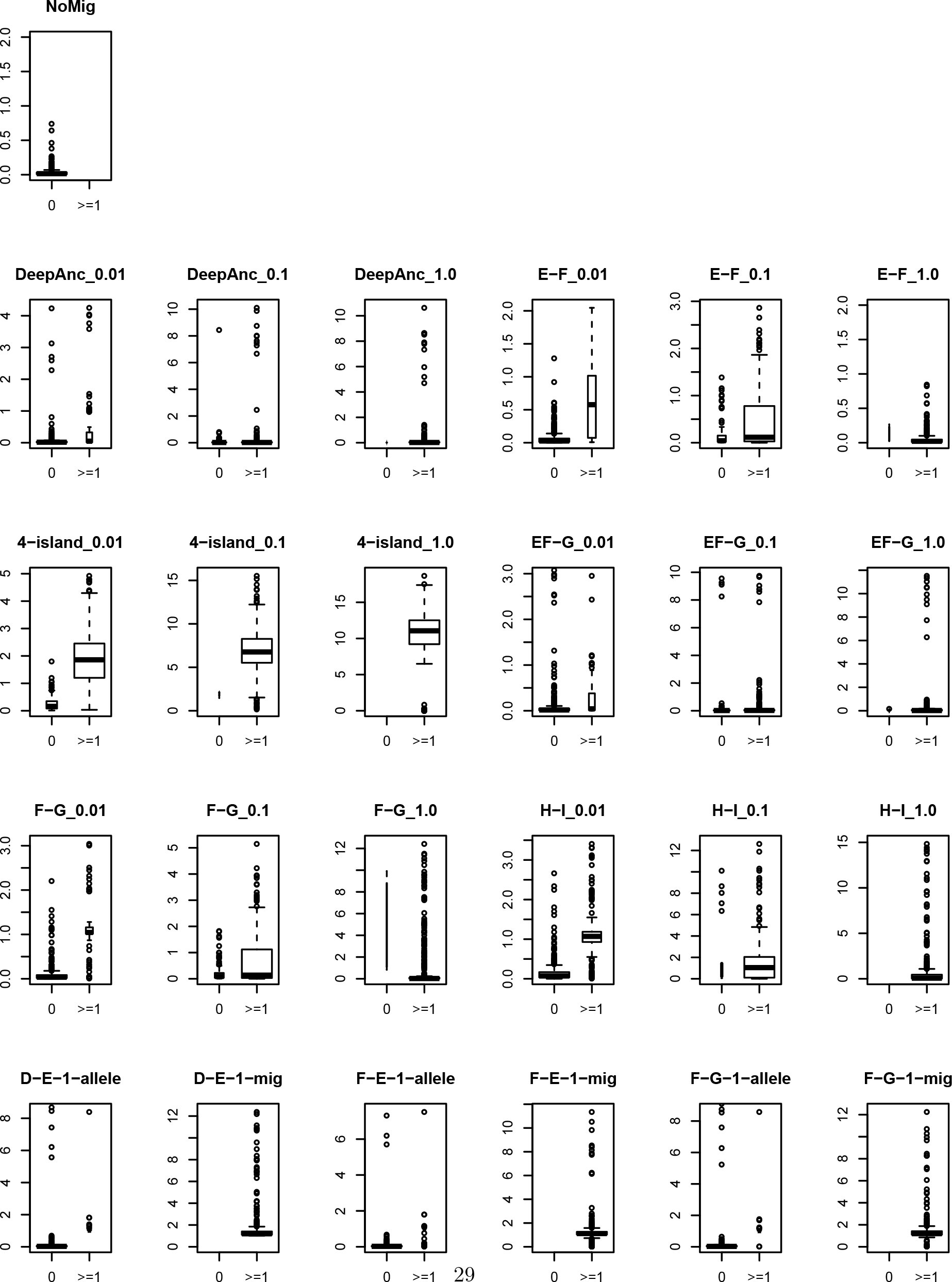
Migration detection for the 10-species scenarios. The scenarios are named as in Figure 6. Other details are as Figure 7.

It is interesting that DENIM identifies TBO64, but not TBO29, as a locus with migration. The posterior mean count of migrations for TBO64 was 1.20, for TBO47 it was 0.47, for TBO26 it was 0.11, and the rest, including TBO29, were well under 0.1. In Belfiore et al. (2008), the individual gene trees were estimated separately, and it appears from their Figure 2 that in TBO64, the relative distance between *Orthogeomys heterodus* and other taxa is considerably smaller than is the case for any other locus.

Other settings were also tried in order assess the sensitivity of the results to the choice of prior. With the simple model, and a prior mean of 0.005 for the migration rate, the method broke down, in a way similar to the tests with no data: the species tree height and migration counts were very large. With the simple model, and a prior mean of 0.0002, the result was similar to Figure 9, although the posterior probability for the ingroup decreased to 0.72. With the flexible model, prior means of 0.001, 0.005, and 0.02 produced similar species trees to Figure 9, although the migration counts and the posterior probabilities for clades were affected.

**Figure 9:**
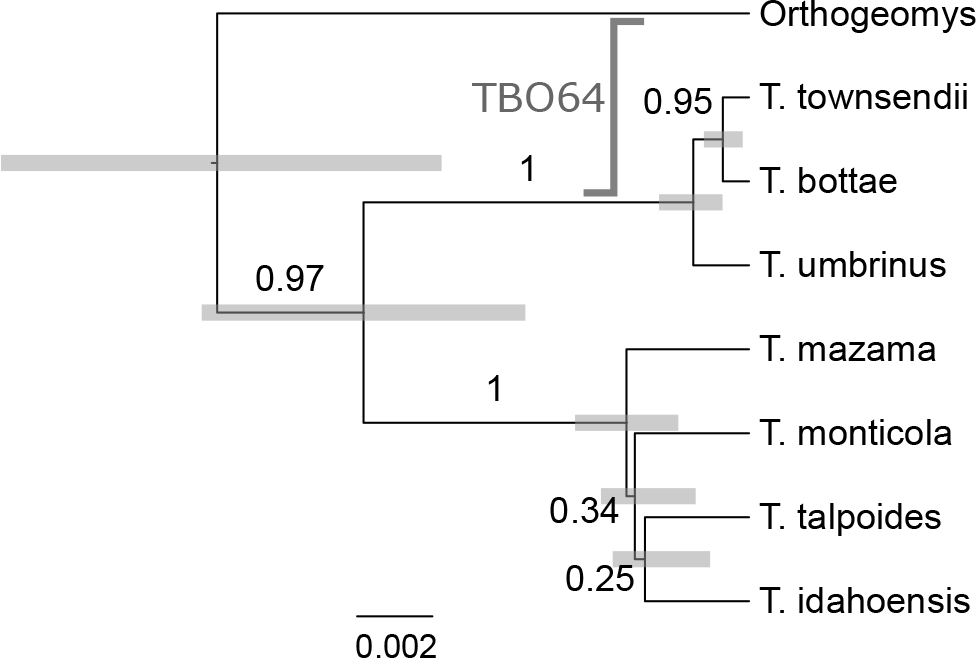
Gopher tree. Posterior clade probabilities are shown next to branches. The node bars are 95% HPDs for the node heights. The migration of of locus TBO64 is also indicated.

## Discussion

If ‘species' are to be defined so that only small amounts of migration can occur between different species, the restriction to only allowing for small migration rates seems reasonable. DENIM can be used to infer a species tree despite a little migration, and can also warn when migration is likely to be present, so that results from other methods which ignore migration can be suitably qualified.

The behaviour of DENIM with no data is far from ideal, and care must be taken to ensure the signal in the data is sufficient to overcome the potential bias from the prior. The embedding scheme means that DENIM can only consider at most one migration per coalescence. The approximation is likely to produce bias if the number of migrations inferred by DENIM is more than a small proportion of the number of coalescences. It is always possible to prevent this by using a prior with a small enough mean, but that may result in bias the other way, inferring too few migrations. With some data sets, it may be that the only conclusion that can be drawn is that there is too little signal and/or too much migration for any sensible estimate of the species tree to be possible using DENIM. If similar results are obtained over a wide range of prior means for the migration rates, the results should be trustworthy.

DENIM combines the tree-island model, which is an elegant mathematical model for speciaton, coalescence, and migration, with a rather crude approximation for sampling the posterior. The two components are quite independent. The partial sampling of the posterior is a trade-off between accuracy on the one hand and computational effort and simplicity of implementation on the other. An exact sampling from the posterior for large data sets when there is a large amount of migration may remain computationally infeasible for many years. However there are almost certainly better compromises to be found than the one currently implemented in DENIM. For example, Palczewski and Beerli (2013) provides an approximation for high rates.

Suppose that all computational problems have been solved. How much data would be needed to get good estimates of the species tree? Hey et al. (2015) shows that good estimates of speciation times can be hard to obtain with small data sets: “for small data sets, with little divergence between samples from two populations, an excellent fit can often be found by a model with a low migration rate and recent splitting time *and* a model with a high migration rate and a deep splitting time.” It may also be that two or more species tree topologies can all achieve excellent fit in models which allow high migration rates, and it would be valuable to find out if this is the case.

It is possible to combine the tree-island model with species delimitation and thus co-estimate the delimitation and the species tree in the presence of migration. The current implementation of DENIM allows this, using the birth-death-collapse model of Jones et al. (2014), but this possibility has not been explored in any detail. The parameter space become even larger, and obtaining useful results in a reasonable amount of time may be very difficult.

DENIM uses the usual birth-death model to provide a prior for the species tree, but this only provides a probability density for the reconstructed tree, with all extinct branches removed. In the presence of migration, there may be gene flow from extinct species which could result in unusually deep coalescences and bias the analysis. DENIM could be extended so that the full tree, including the extinct branches is sampled from. There is no upper limit to the number of extinct branches that could exist, so again there are more computational difficulties. In principle, this could allow the detection of some extinct species from genetic data alone.

## Acknowledgments

I thank Adam Leaché for supplying the MCcoal control files used to generate the simulated data.

## ONLINE APPENDIX: Supplementary figures

**Figure 10:**
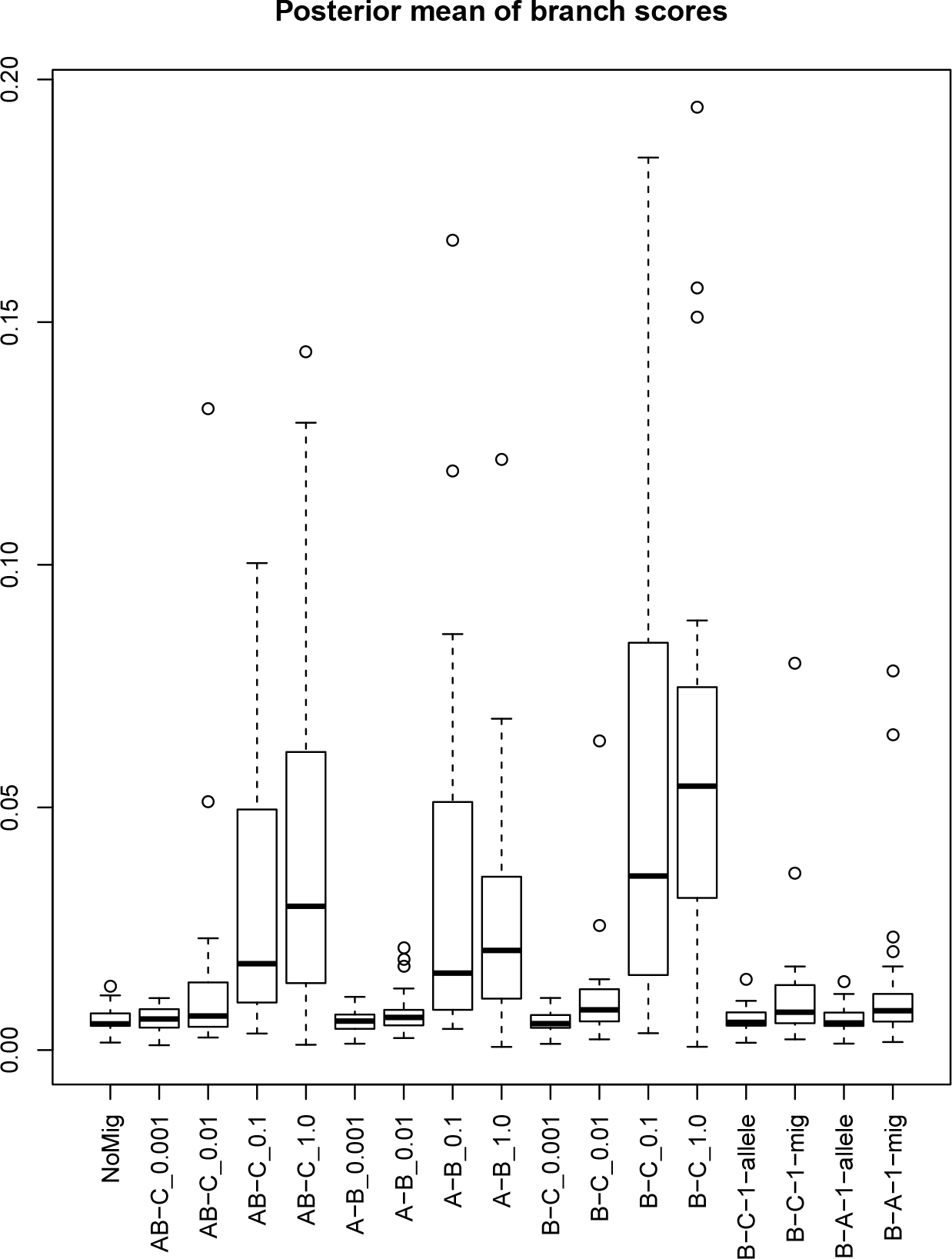
Branch scores for the 4-species scenarios, based on 100 replicates. “NoMig” is the scenario with no migration. The other names describe the pairs of species tree branches which have migration, followed by the migration rate, or “allele” meaning a single locus introgression, or “mig”, meaning a single migrant. The flexible model was used with sa prior mean of 0.00125.

**Figure 11:**
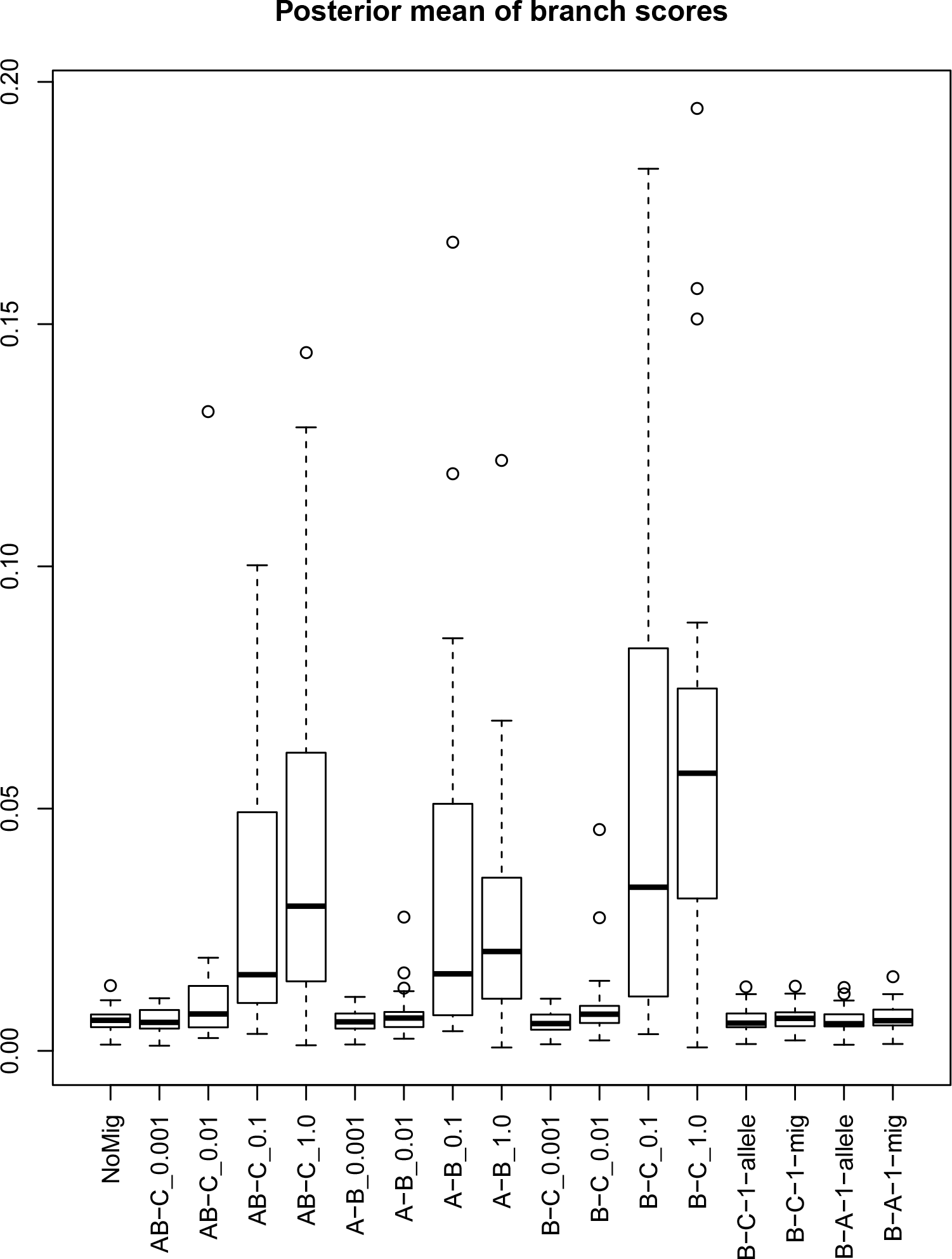
As Figure 10 except that the prior mean was 0.005.

**Figure 12:**
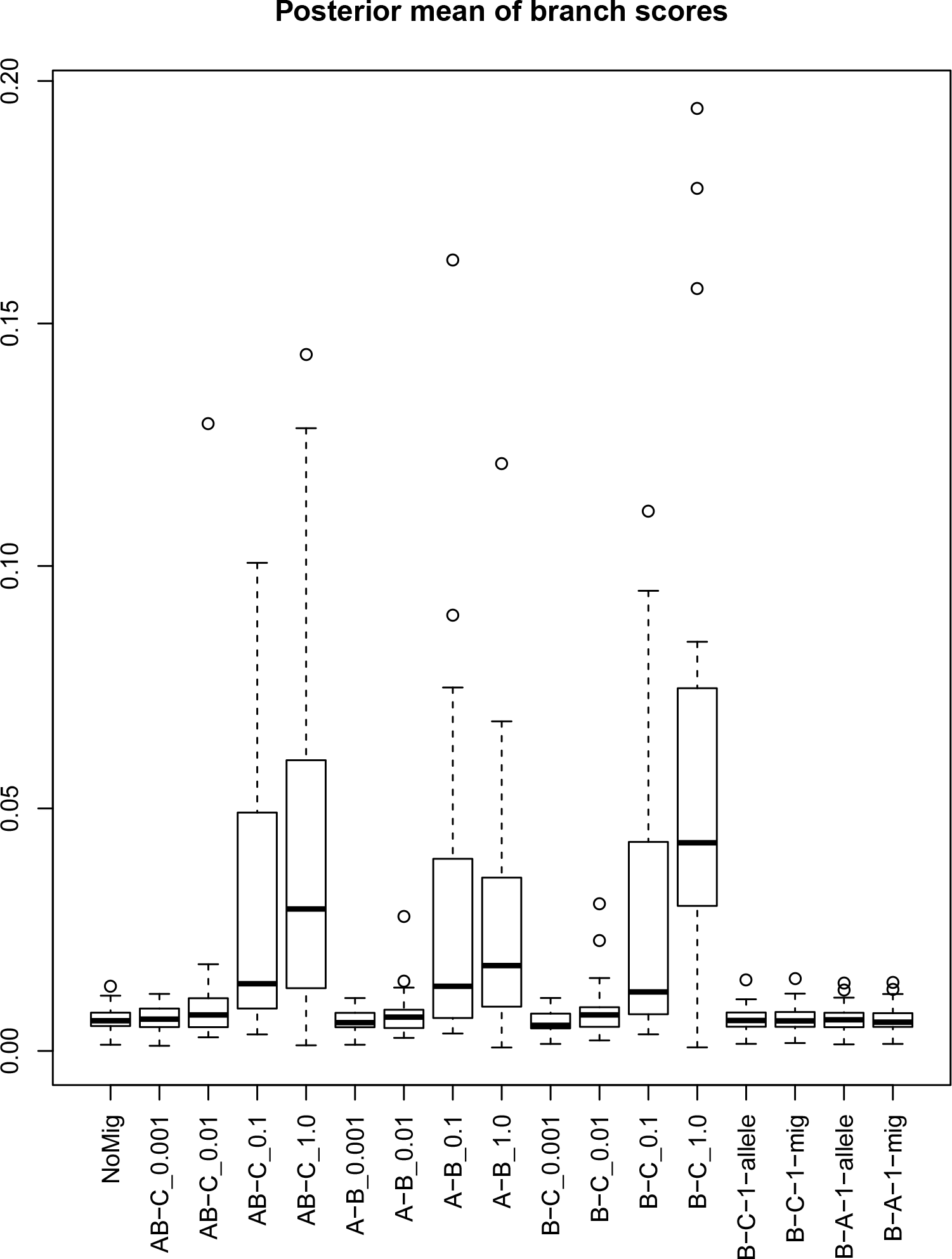
As Figure 10 except that the prior mean was 0.02.

**Figure 13:**
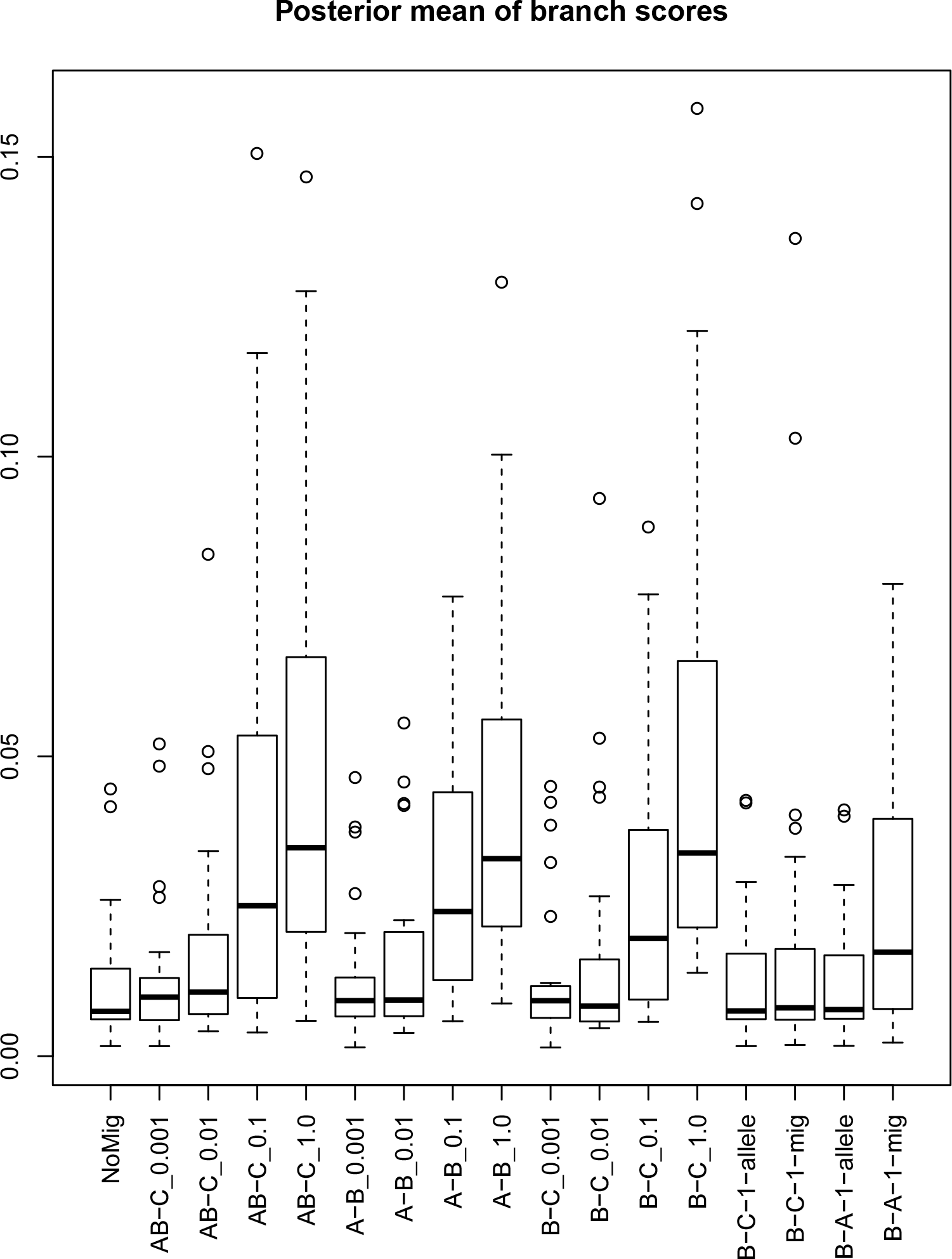
As Figure 10 except that the prior mean was 0.08.

**Figure 14:**
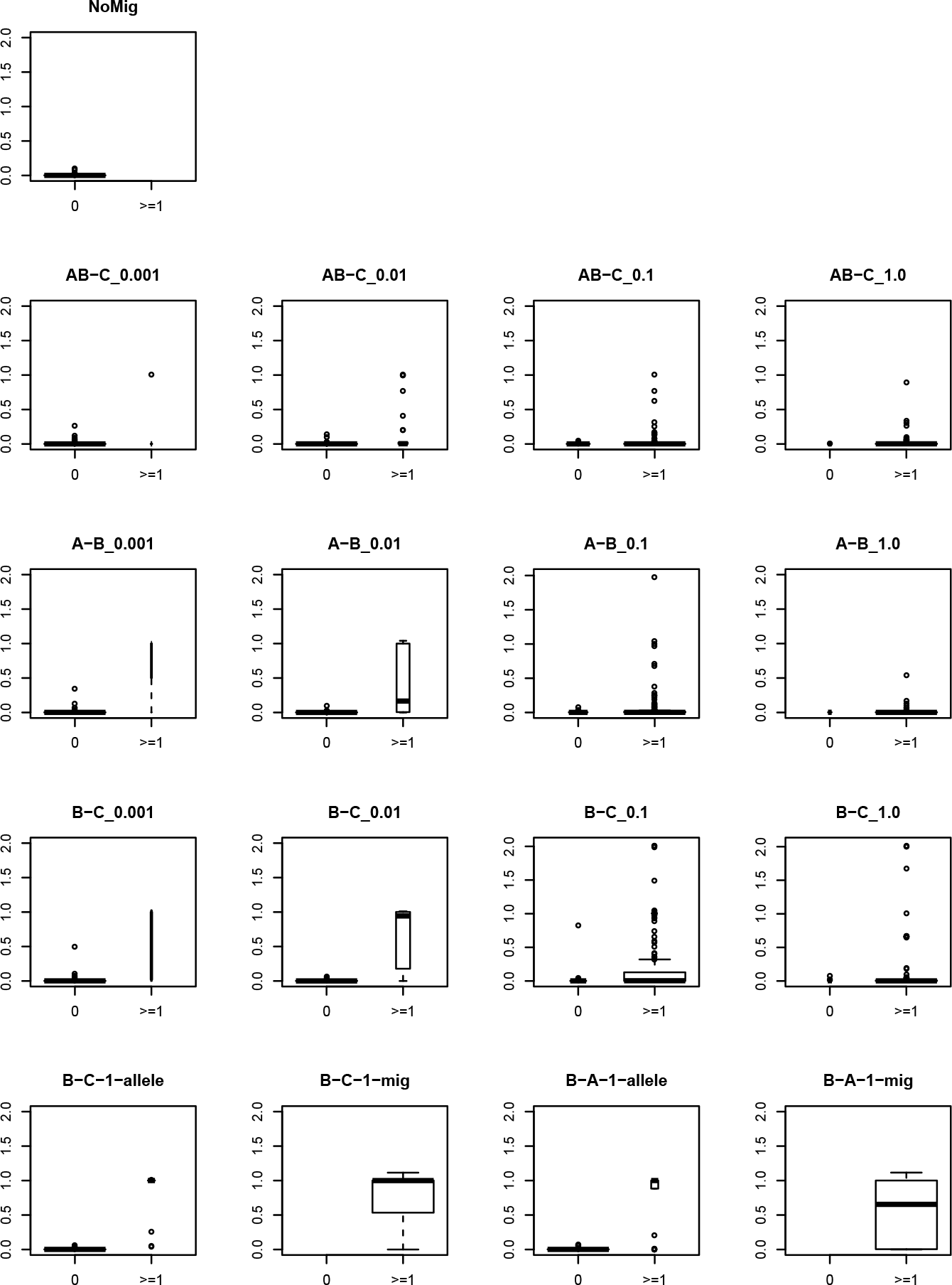
Migration detection for the 4-species scenarios. Each boxplot show the posterior mean count of migrations for the two cases that migration is present or absent in a locus in the simulated data. The width of the boxes is proportional to the number of cases. The flexible model was used with a prior mean of 0.00125.

**Figure 15:**
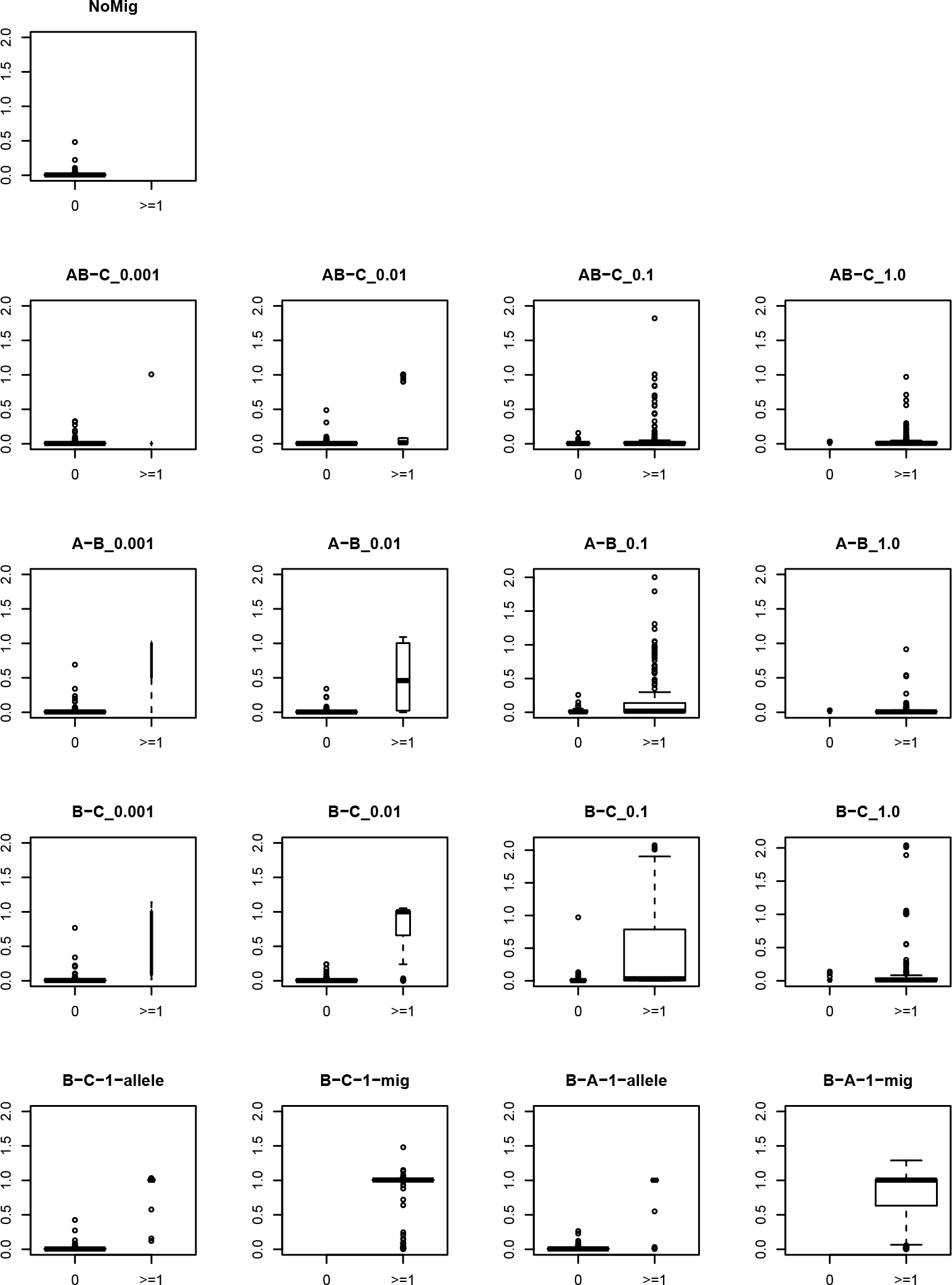
As Figure 14 except that the prior mean was 0.005.

**Figure 16:**
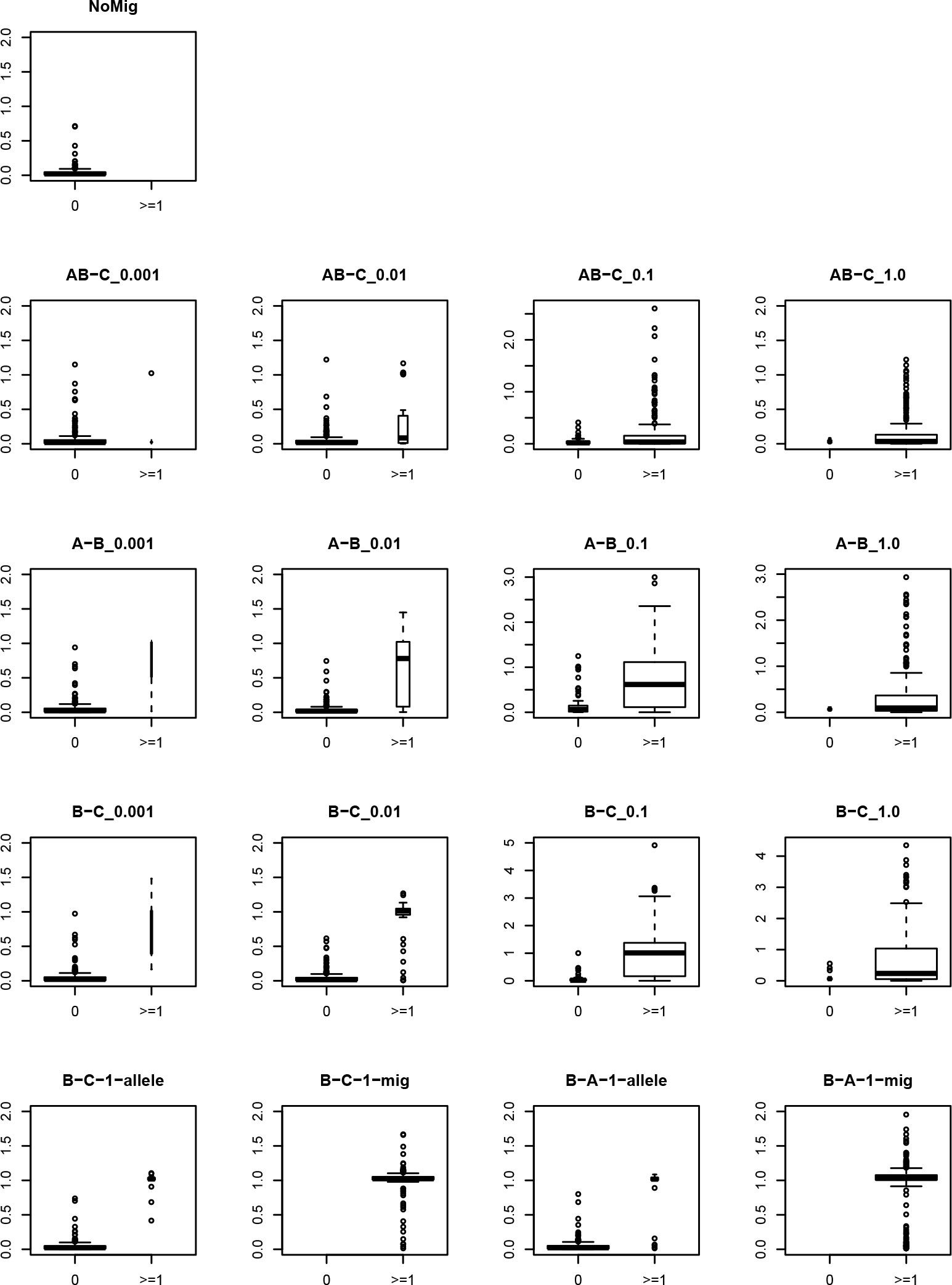
As Figure 14 except that the prior mean was 0.02.

**Figure 17:**
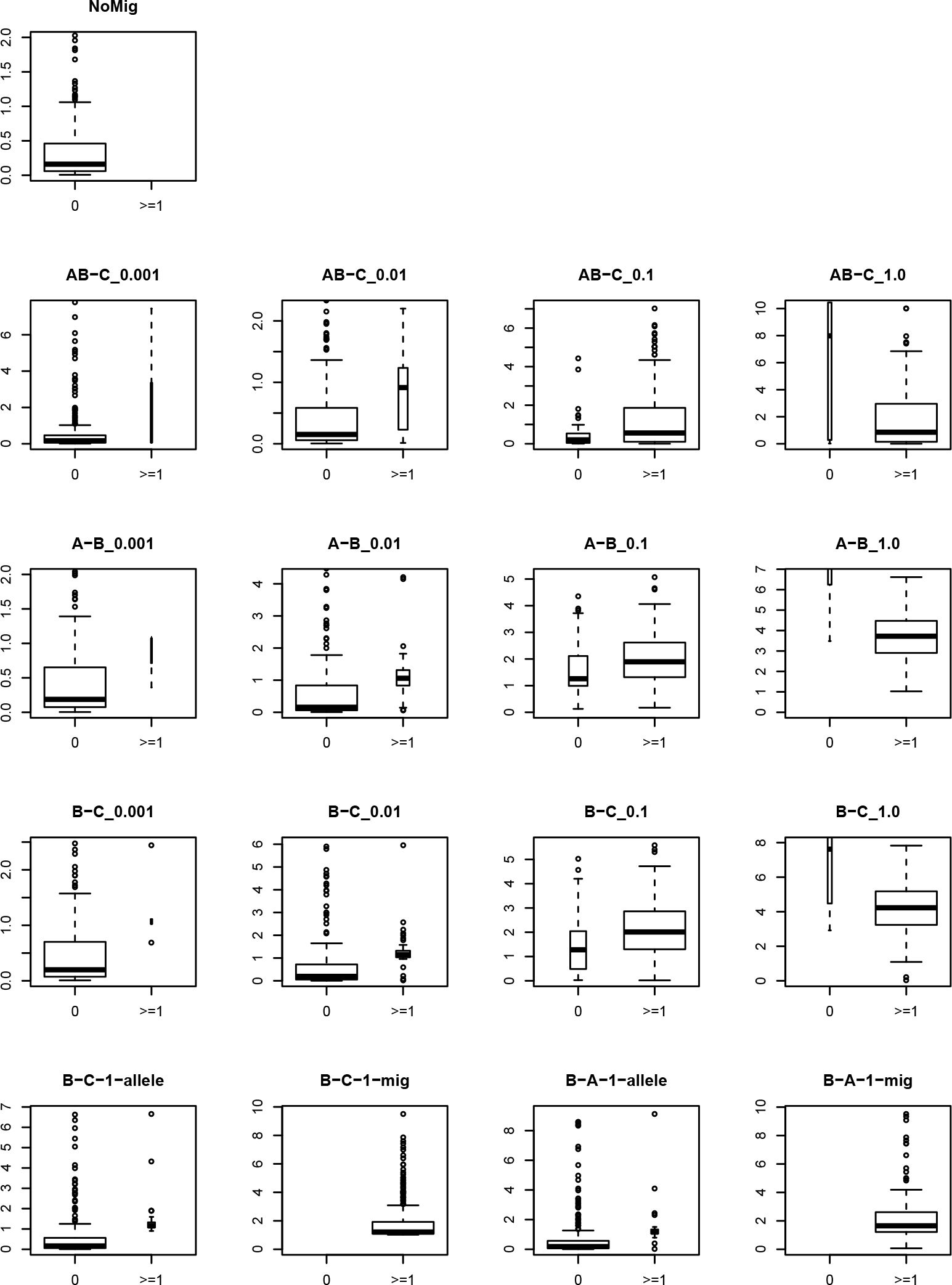
As Figure 14 except that the prior mean was 0.08.

## References

Peter Beerli and Joseph Felsenstein. Maximum likelihood estimation of a migration matrix and effective population sizes in n subpopulations by using a coalescent approach. Proceedings of the National Academy of Sciences, 98(8):4563–4568, 2001.

Natalia M. Belfiore, Liang Liu, and Craig Moritz. Multilocus phylogenetics of a rapid radiation in the genus thomomys (rodentia: Geomyidae). Systematic Biology, 57(2):294, 2008. doi:10.1080/10635150802044011. URL + http://dx.doi.org/10.1080/10635150802044011.

Daniel A Dalquen, Tianqi Zhu, and Ziheng Yang. Maximum likelihood implementation of an isolation-with-migration model for three species. Systematic Biology, 00:00–00, 2016.

Greg Ewing and Rodrigo Allen. Estimating population parameters using the structured serial coalescent with Bayesian MCMC inference when some demes are hidden. Evolutionary Bioinformatics, 2, 2006.

Henrique V. Figueiró, Gang Li, Fernanda J. Trindade, Juliana Assis, Fabiano Pais, Gabriel Fernandes, Sarah H. D. Santos, Graham M. Hughes, Aleksey Komissarov, Agostinho Antunes, Cristine S. Trinca, Maíra R. Rodrigues, Tyler Linderoth, Ke Bi, Leandro Silveira, Fernando C. C. Azevedo, Daniel Kantek, Emiliano Ramalho, Ricardo A. Brassaloti, Priscilla M. S. Villela, Adauto L. V. Nunes, Rodrigo H. F. Teixeira, Ronaldo G. Morato, Damian Loska, Patricia Saragüeta, Toni Gabaldón, Emma C. Teeling, Stephen J. O’Brien, Rasmus Nielsen, Luiz L. Coutinho, Guilherme Oliveira, William J. Murphy, and Eduardo Eizirik. Genome-wide signatures of complex introgression and adaptive evolution in the big cats. Science Advances, 3(7), 2017. doi:10.1126/sciadv.1700299. URL http://advances.sciencemag.org/content/3/7/e1700299.

J Heled and A Drummond. Bayesian inference of species trees from multilocus data. Mol. Biol. Evol., 27: 570–580, 2010.

J Hey. Isolation with migration models for more than two populations. Mol Biol Evol, 27:905–920, 2010.

J Hey and R Nielsen. Multilocus methods for estimating population sizes, migration rates and divergence time, with applications to the divergence of drosophila pseudoobscura and d. persimilis. Genetics, 167: 747–760, 2004.

J Hey and R Nielsen. Integration within the Felsenstein equation for improved Markov chain Monte Carlo methods in population genetics. PNAS, 104:2785–2790, 2007.

Jody Hey, Yujin Chung, and Arun Sethuraman. On the occurrence of false positives in tests of migration under an isolation-with-migration model. Molecular Ecology, 24(20):5078–5083, 2015. ISSN 1365-294X. doi:10.1111/mec.13381. URL http://dx.doi.org/10.1111/mec.13381.

Richard R Hudson et al. Gene genealogies and the coalescent process. Oxford surveys in evolutionary biology, 7(1):44, 1990.

Nathon D. Jackson, Ariadna E. Morales, Bryan C. Carstens, and Brian C. OMeara. Phrapl: Phylogeographic inference using approximate likelihoods. Systematic Biology, 66(6):1045–1053, 2017. doi:10.1093/sys-bio/syx001. URL + http://dx.doi.org/10.1093/sysbio/syx001.

Graham Jones. Algorithmic improvements to species delimitation and phylogeny estimation under the multispecies coalescent. Journal of Mathematical Biology, 2016. doi:10.1007/s00285-016-1034-0. URL http://link.springer.com/article/10.1007/s00285-016-1034-0.

Graham Jones, Zeynep Aydin, and Bengt Oxelman. DISSECT: an assignment-free Bayesian discovery method for species delimitation under the multispecies coalescent. Bioinformatics, 2014. doi:10.1093/bioinformatics/btu770.

J.F.C. Kingman. The coalescent. Stochastic processes and their applications, 13(3):235–248, 1982.

A D Leaché, R B Harris, B Rannala, and Z Yang. The influence of gene flow on species tree estimation: A simulation study. Systematic Biology, 63(1):17–30, 2014.

S. H. Martin, K. K. Dasmahapatra, N. J. Nadeau, C. Salazar, J. R. Walters, F. Simpson, and C. D. Jiggins. Genome-wide evidence for speciation with gene flow in heliconius butterflies. Genome Research, 23(11): 1817–1828, 2013.

Patrik Nosil. Speciation with gene flow could be common. Molecular Ecology, 17(9):2103–2106, 2008. ISSN 1365-294X. doi:10.1111/j.1365-294X.2008.03715.x. URL http://advances.sciencemag.org/content/http://dx.doi.org/10.1111/j.1365-294X.2008.03715.x.

Huw A. Ogilvie, Remco R. Bouckaert, and Alexei J. Drummond. Starbeast2 brings faster species tree inference and accurate estimates of substitution rates. Mol Biol Evol, 2017. doi:10.1093/molbev/msx126.

Michal Palczewski and Peter Beerli. A continuous method for gene flow. Genetics, 194(3):687–696, 2013. ISSN 0016-6731. doi:10.1534/genetics.113.150904. URL http://www.genetics.org/content/194/3Z687.

Martyn Plummer, Nicky Best, Kate Cowles, and Karen Vines. CODA: Convergence diagnosis and output analysis for MCMC. R News, 6(1):7–11, 2006. URL http://CRAN.R-project.org/doc/Rnews/.

Frank E. Rheindt, Matthew K. Fujita, Peter R. Wilton, and Scott V. Edwards. Introgression and phenotypic assimilation in zimmerius flycatchers (tyrannidae): Population genetic and phylogenetic inferences from genome-wide snps. Systematic Biology, 63(2):134–152, 2014. doi:10.1093/sysbio/syt070. URL + http://dx.doi.org/10.1093/sysbio/syt070.

Konstantin Romaschenko, Nuria Garcia-Jacas, Paul M. Peterson, Robert J. Soreng, Roser Vilatersana, and Alfonso Susanna. Miocenepliocene speciation, introgression, and migration of patis and ptilagrostis (poaceae:Stipeae). Molecular Phylogenetics and Evolution, 70:244–259, 2014. ISSN 1055-7903. doi: http://dx.doi.org/10.1016/j.ympev.2013.09.018. URL http://www.sciencedirect.com/science/article/pii/S1055790313003734.

Claudia Solís-Lemus and Cécile Ané. Inferring phylogenetic networks with maximum pseudolikelihood under incomplete lineage sorting. PLOS Genetics, 12(3):1–21, 03 2016. doi:10.1371/journal.pgen.1005896. URL https://doi.org/10.1371/journal.pgen.1005896.

Yuan Tian and Laura S Kubatko. Distribution of coalescent histories under the coalescent model with gene flow. Molecular Phylogenetics and Evolution, 105:177–192, 2016.

D Wen, Y Yu, and L Nakhleh. Bayesian inference of reticulate phylogenies under the multispecies network coalescent. PLoS Genet, 12(5):e1006006, 2016. URL https://doi.org/10.1371/journal.pgen.1006006.

Ziheng Yang. The BPP program for species tree estimation and species delimitation. Current Zoology, 61 (5):854–865, 2015.

Chi Zhang, Huw A Ogilvie, Alexei J Drummond, and Tanja Stadler. Bayesian inference of species networks from multilocus sequence data. bioRxiv, 2017. doi:10.1101/124982. URL http://www.biorxiv.org/content/early/2017/04/06/124982.

